# Early skin seeding regulatory T cells modulate PPARγ-dependent skin pigmentation

**DOI:** 10.1101/2023.10.17.561843

**Authors:** Inchul Cho, Jessie Z. Xu, Hafsah Aziz, Prudence PokWai Lui, Boyu Xie, Pei-Hsun Tsai, Hee-Yeon Jeon, Jinwook Choi, Shahnawaz Ali, Niwa Ali

## Abstract

The maintenance of adult tissue homeostasis is dependent on the functional cross-talk between stem cells (SCs) and tissue-resident immune cells. This reciprocal relationship is also essential for tissue organogenesis during early life. The skin harbors a relatively large population of Regulatory T cells (Tregs) that accumulate within the first two weeks after birth. A functional role for early skin seeding Tregs (ETregs) during the first week of life is currently unexplored. Here, we show that skin Tregs are detected early as postnatal day 3 (P3) where they localize to hair follicle (HF) structures and enter a dynamic flux of activation marker expression. Punctual ETreg depletion from P6-P8, but not later, results in defective HF melanocyte SC (MeSC) mediated skin pigmentation in juvenile life. Transcriptomic analysis of the whole skin on P9 exhibited immediate and pronounced changes in MeSC markers and perturbation of Peroxisome proliferator-activated receptor-γ (PPARγ) target genes in the HF. Accordingly, punctual ETreg depletion combined with short-term PPARγ agonization restored skin pigmentation. Single cell profiling of P9 skin revealed that PPARγ signalling activity is preferentially diminished in the HF epithelium upon loss of ETregs. Finally, we explored changes in the single cell transcriptome of the human tissue disorder, vitiligo, characterized by a lack of melanin and consequent skin depigmentation. These analyses showed that the HF cells from lesional vitiligo skin exhibited a significant downregulation in PPARγ pathway activation, relative to heathy controls. Overall, ETregs in neonatal skin are critical for sustaining HF PPARγ signaling, which is vital for facilitating MeSC mediated skin pigmentation during postnatal development.

**One Sentence Summary:** PPARγ pathway functions downstream of neonatal Tregs to regulate melanocyte stem cell function.

## INTRODUCTION

Tissue organogenesis during neonatal life is facilitated by temporarily and spatially coordinated crosstalk between immune cells and stem cells (SCs) in the tissue microenvironment. Barrier sites such as the skin harbor relatively large proportions of the CD4 lineage of Foxp3+ regulatory T cells (Tregs) that reside proximal to distinct adnexal structures, such as hair follicle (HFs) (*1–3*). The HFs represent an important immune cell niche in skin, but also accommodate distinct populations of SCs. These include HFSCs and melanocyte SCs (MeSCs), governing hair shaft production and skin pigmentation, respectively.

An intimate association between skin Tregs and the HF epithelia has been well established in recent years. During development, Treg seed the skin during the first week of life in conjunction with HF morphogenesis, peaking on postnatal day 13 (P13) (*4*). Tregs resident in P8-15 skin are essential for establishing both tolerance to commensal bacterial antigens and suppression of type 2 helper T cell mediated fibrous pathology (*5*, *6*). In adult skin, Tregs expressing the Notch ligand Jagged-1, are required for hair regeneration via promotion of HFSC differentiation. This SC regulatory function of Tregs is orchestrated largely independently of canonical immunosuppressive pathways. Conversely, during subacute skin injury, Treg control of inflammatory Th17 responses facilitates HFSC mediated barrier regeneration. While these studies have ascribed distinct and specialised roles for adult Tregs and neonatal Tregs during the second week of life (*7*), a functional role for the foremost afferent population of skin seeding Tregs during the first week of life has not been explored. Specifically, the existence of a neonatal Treg-SC axis during this specific window of time and its relevance to the homeostatic development and function of the skin remains unknown.

Here, we demonstrate that neonatal Tregs in P6-P8 skin are highly activated and preferentially reside in close association with MeSCs. Transient loss of Tregs during this time window, but not later, results in defective melanocyte function *in vivo*, as evidenced by disrupted skin pigmentation in later life. Notably, the transcriptomic changes in melanocytes precede the inflammatory responses induced by Treg depletion, suggesting Tregs may regulate MeSCs independently of secondary mediators such as CD4^+^ FoxP3^-^ effector T cells (Teffs) or CD8^+^ cytotoxic T cells. Furthermore, we observed an immediate dysregulation of transcripts associated with the Peroxisome proliferator-activated receptor (PPAR) signalling pathway. Additionally, we identify changes in PPARγ signalling activity throughout developmental stages of human skin and in the pigment-deficient disease state of vitiligo. We define neonatal Treg regulation of the PPARγ pathway as a mechanistic link between early skin development and functional MeSC melanogenic activity for establishing pigmentation of skin. Taken together, our findings reveal that neonatal Tregs during the first week of life are essential for the establishment of SC function during postnatal development of the skin.

## RESULTS

### Skin Tregs are highly activated on Postnatal day 6

The accumulation of Tregs in skin peaks on postnatal day 13 (P13) of life, likely derived from lymphocytic seeding from lymphoid organs (*6*). At this time-point Tregs display a highly proliferative and activated profile. We sought to perform immune profiling at earlier timepoints, prior to P13, upon first entry of Tregs in skin. We first characterized Treg abundance and phenotypic marker expression in steady-state C57BL/6 neonatal skin and skin-draining lymph node (SDLNs) on P6, P9, P12, and later P28 (juvenile) and P49 (adult) timepoints.

Flow cytometric profiling revealed the presence of Tregs at detectable but low numbers on P3 (**Figure 1A-B**; full gating strategy is shown in **Supplementary figure 1**). The abundance and proportion of CD4^+^Foxp3^+^ Tregs steadily increased and peaked during the second week after birth, which is consistent with previous studies (**Figure 1A-B**) (*6*). This wave of Treg accumulation in P12 skin was unique to Tregs as our global analysis of other major skin-resident T cell subsets during this time frame did not display a similar pattern, namely Foxp3^-^ T effector cells (Teffs) and CD8^+^ T cells (**Supplementary figure 2A-F**). Of note, Treg percentages on P8 and P12 were comparable to that found in adult skin on P49 (**Figure 1A**), corroborating a recent study reporting similar findings (*8*). By contrast, the percentage of FoxP3^+^ Tregs in SDLNs was less variable and remained steady throughout the analysis period. The proliferative index of Teffs, CD8+ T cells, and Tregs was the highest in P6 skin but similar amongst all T cell subsets in the three time-points assessed, as evidenced by equivalent levels of Ki67 expression. However, in P9 SDLNs, Tregs were more proliferative relative to other T cell subsets (**Supplementary figure 2C, 2F**).

**Figure 1.**
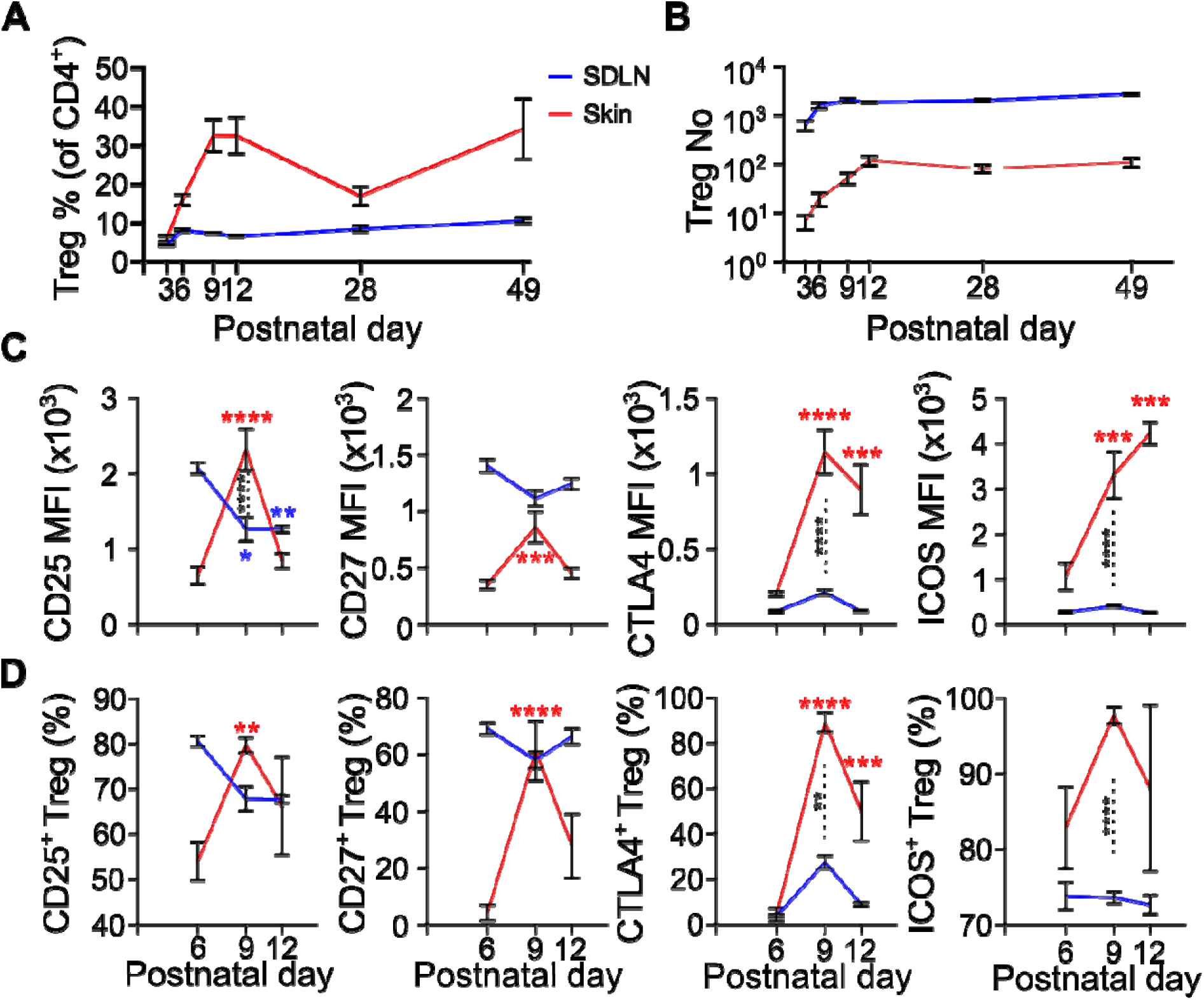
Neonatal skin Tregs accumulate during the first 12 days of life and undergo phenotypic changes. **A-D)** Flow cytometric profiling of skin CD4^+^ FoxP3^+^ regulatory T cells (Tregs) (n=4-6 biological replicates) pooled from 2 independent experiments. **A)** Percentage of Tregs in the skin (of total CD4^+^ T cells) in the skin and the skin-draining lymph node (SDLN) on postnatal day 3 (P3), P6, P9, P12, P28 and P49. **B)** Absolute number of Tregs in the skin (per 6 cm^2^ of skin) and SDLN (per 10^5^ of total cells) on P6, P9 and P12. **C)** Median fluorescence intensities (MFIs) Treg phenotypic markers, CD25, CD27, ICOS and CTLA4. **D)** Percentage of Tregs expressing CD25, CD27, ICOS and CTLA4 (of total Tregs). Graphs show mean ± S.E.M. (n=4-6 biological replicates). P value was calculated using One way ANOVA against P6. Two way ANOVA with Sidak’s multiple comparison test was used to test skin Tregs against SDLN Tregs. ****p<0.0001, ***p<0.001, **p<0.01, *p<0.05, ns p>0.05.

Given the rapid influx of Tregs in skin during the early neonatal period of P6-P12, we assessed the expression of the activation-associated markers CD25, CD27, CTLA4, and ICOS. Between P6 to P9, the median flourescence intensity (MFI) values of all markers, except CD27, were significantly higher in P9 skin Tregs relative to their SDLN counterparts (**Figure 1C**). In addition, the MFI of all markers were upregulated by at least 2-fold between P6 and P9 skin Tregs, but not in SDLN Tregs (**Figure 1C**). The proportions of Treg activation marker expression largely resembled the fluctuations observed in MFI values for both skin and SDLN compartments (**Figure 1D**). Interestingly, later skin seeding Tregs (LTregs) in P12 skin expressed lower levels of CD25, CD27, and CTLA4 relative to P9. Taken together, these data suggest P3-P12 is an important interval where Tregs accumulate in skin and identifies P6-P9 as a specific time window where early skin seeding Tregs (ETregs) become highly, but transiently, activated in the tissue.

### Neonatal Tregs suppress inflammation and skin pigmentation in later life

Given that Treg activation and proliferation is highly dynamic in P6-P12 skin, we hypothesized that Tregs in this time window may play an important role in regulating either inflammatory responses or postnatal skin development, or both. To test this, we utilized *Foxp3-DTR* transgenic mice where the diphtheria toxin receptor (DTR) is expressed ahead of the *Foxp3* promoter (*9*). Following administration of DT, these mice permit highly robust Treg depletion in both SDLNs and skin (*5*, *10*, *11*). To ablate Tregs from P6-P12, we administered DT on P6, P8, P10 and P12 (hereafter referred to as the “ΔΤreg” group). We chose P28 as the timepoint for analysis as the major hallmarks of stem cell (SC) mediated skin development are manifested at this age, namely hair follicle SC (HFSC) activity and melanocyte SC (MeSC) mediated skin pigmentation.

While Treg sufficient controls on P28 developed normal appearing dorsal skin with black pigmentation, this was markedly impacted in ΔTreg animals, where the skin failed to pigment (**Supplementary figure 3A-B**). This phenotype was confirmed histologically using Fontana & Masson (F&M) staining that detected melanin granules produced by MeSCs in Treg sufficient controls but not in ΔΤreg skin (**Supplementary figure 3C**). To ascertain whether stem cell regulation is impacted upon Treg depletion, we profiled CD34^+^ Itga6^+^ HFSCs and CD117^+^ MeSCs by flow cytometry (**Supplementary figure 3D-E**). Both the proliferation and abundance of HFSCs were unaffected in ΔTreg skin relative to controls on P28. However, MeSC proliferation was markedly reduced in ΔTreg skin. These findings suggest the melanogenic function of MeSCs in the HFs is under Treg control during the P6-P12 time frame, whereas HFSCs are not impacted.

To determine if loss of neonatal Tregs results in systemic inflammation in later life we firstly monitored body weight gain. Depletion of Tregs during this six-day window resulted in weight reduction by P28. This was despite of a repopulating Treg presence that was significantly higher than Treg-sufficient controls (**Supplementary figure 3F**). In addition, immune profiling of all major skin-resident T cell subsets revealed an increased abundance and activation status of Teffs and CD8^+^ T cells on P13 – only 1 day after the Treg depletion regimen (**Supplementary figure 3G-H**). This inflammatory phenotype persisted even at the later timepoint of P28 (**Supplementary figure 3I-K**). In particular, CD8^+^ T cells were highly proliferative and outnumbered Tregs, as evidenced by an increased CD8:Treg ratio in ΔTreg mice relative to controls. These data suggest there is an immediate inflammatory response following the six-day Treg depletion regime that may persist until later life. Indeed, the same Treg depletion regimen in adult mice does not cause overt skin inflammation (*10*), supporting the idea that neonatal and adult Tregs are functionally distinct (*7*). Overall, the loss of neonatal Tregs from P6-P12 leads to defective MeSC mediated melanogenesis and a consequent defect in skin pigmentation, that may be associated with a systemic inflammatory response.

### Early skin seeding Tregs (ETregs) are required for skin pigmentation

In our initial experiments, we chose a 4-dose DT administration to deplete Tregs from P6-P12 **(Supplementary figure 3).** While this resulted in a defective skin pigmentation phenotype, it was also accompanied by reduced weight gain and an immediate inflammatory response that persisted until P28 (**Supplementary figure 3F-K**). Thus, we set out to determine whether defective MeSC function was a result of prolonged systemic inflammation and to also define a precise ‘window’ of time for Treg requirement. Given ETregs acquire a highly activated profile between P6 to P9, and the less activated profile of LTregs in P12 skin (**Figure 1C-D**), we sought to address how transient loss of Tregs at these two timepoints would impact both skin development and local inflammation. To do so, we implemented a 2-dose DT regimen by administering DT on P6 and P8 to deplete ETregs (hereafter referred to as the “ΔETreg” group), and on P10 and P12 to deplete LTregs (hereafter referred to as the “ΔLTreg” group). Efficient Treg depletion in the skin was confirmed one day after the last DT injection on P9 (for the ETreg group) and P13 (for LTreg group) (**Supplementary figure 4A-B**). Notably, transient depletion of Tregs in the ETreg group recapitulated the attenuation of dorsal skin pigmentation on P28 observed in the 4-dose DT-treated ΔTreg group (**Figure 2A-E**). In contrast, the skin of ΔLTreg animals was not affected relative to controls (**Figure 2B-E**). Histologic F&M examination showed a significant reduction in melanin production in ΔETreg, but not in ΔLTreg animals, as quantified using a skin pigmentation index (**Figure 2D-E**). Importantly, Treg ablation during the P6 to P8 window in ΔETreg animals had no significant effect on weight gain relative to Treg sufficient control groups (**Supplementary figure 4C**). These results indicate an early 2-dose depletion regimen targeting ETregs, but not LTregs, impairs skin pigmentation without compromising animal fitness in later life.

**Figure 2.**
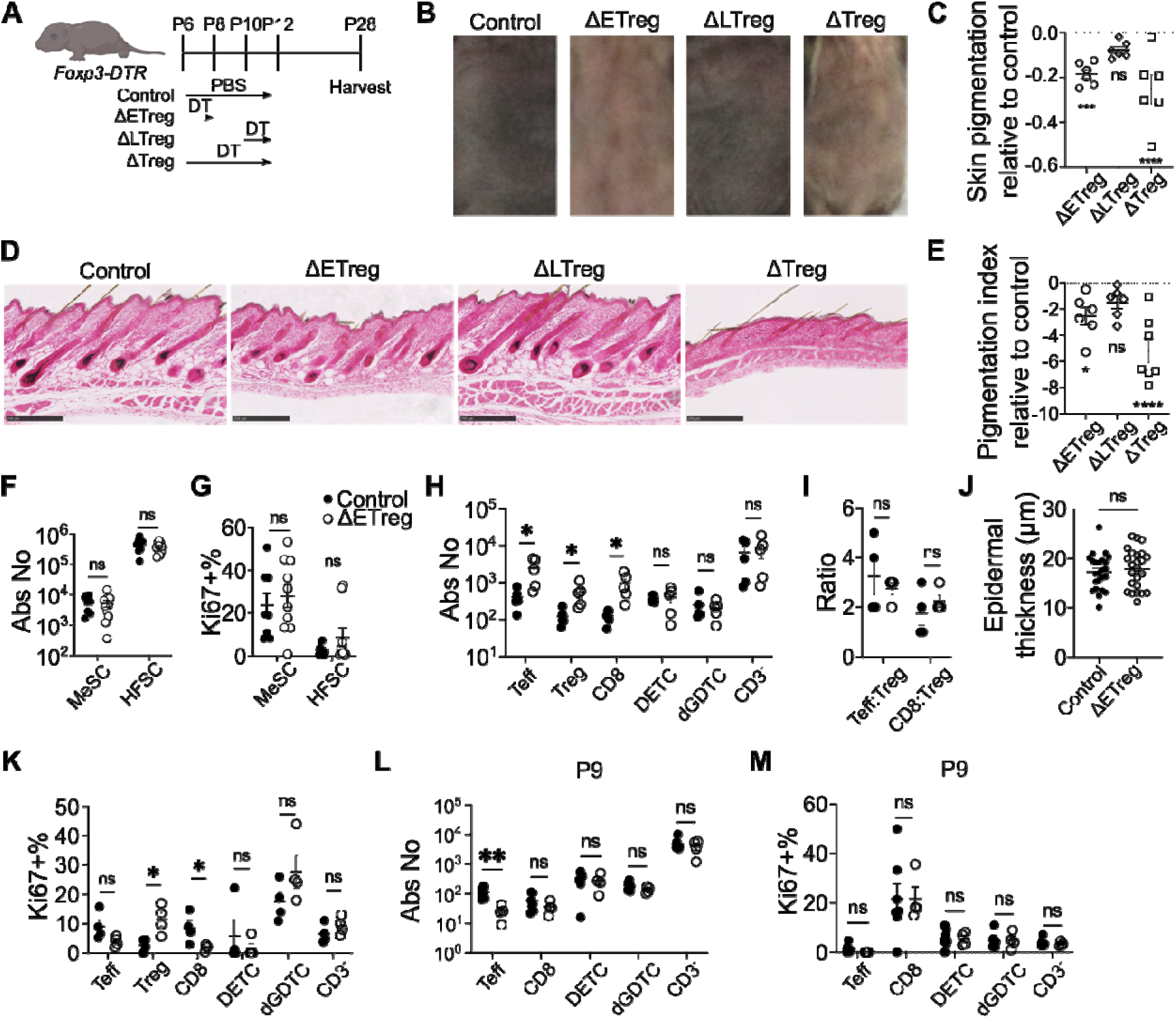
Early neonatal Tregs are required for skin pigmentation. **A)** Schematic outline of experimental timeline. *Foxp3-DTR* transgenic mice received PBS on postnatal day 6 (P6), P8, P10 and P12 (Control), DT on P6 and P8 (ΔETreg), on P10 and P12 (ΔLTreg), and on P6, P8, P10 and P12 (ΔTreg). Skin tissues were harvested on P28. **B)** Representative Fontana & Masson staining, showing black melanin pigments. Pigmentation area was quantified and normalised against PBS control group. Scale bars represent 250 µm. **C)** Quantification of pigmentation index normalised to PBS-injected controls. **D-E)** Flow cytometric quantification of melanocyte stem cell (MeSC) and hair follicle stem cell (HFSC) **D)** Abundance and **E)** Proliferating Ki67^+^ cells. **F-I)** Flow cytometric characterisation of the skin on P28. **F)** Absolute number of skin-resident immune cells. **G)** Proliferating Ki67^+^ immune cells. **H)** CD4^+^ FoxP3^-^ effector T cell (Teff):CD4^+^ FoxP3^+^ Treg ratio and CD8^+^ T cell (CD8):Treg ratio. **I)** Median fluorescence intensity of Treg phenotypic markers, CD25, CD27, ICOS and CTLA4. Cells were pre-gated as CD4^+^ FoxP3^+^ Tregs. **J)** Histological quantification of epidermal thickness. **K-L)** Flow cytometri characterisation of skin-resident immune cells on P9. **K**) Absolute number. **L)** Ki67^+^ immune cells. Graph show mean ± S.E.M. **B-J)** Data is representative of 4 independent experiments. **K-L)** Data are representative of two independent experiments. Unpaired t-test. ****p<0.0001, **p<0.01, *p<0.05, n p>0.05.

Next, we performed flow cytometry to test the hypothesis that defective skin pigmentation is linked to a disruption of SC maintenance and/or activation. Profiling of MeSCs and HFSCs showed that both the abundance and proportion of Ki67 expression remains comparable between control and ΔETreg groups on P28 (**Figure 2F-G**). Therefore, SC maintenance is unaffected in the absence of ETregs.

Given the important role of T cells in disorders of skin pigmentation (*12*), we assessed whether T cell numbers are impacted in ΔETreg skin on P28. Indeed, the skin T cell pool expands, but maintain homeostatic levels of proliferation, Teff:Treg ratios, and CD8:Treg cell ratios (**Figure 2H-K**). We also assessed for the hallmark indicator of global tissue inflammation by quantifying epidermal hyperproliferation of the skin. Histological quantification of epidermal thickness showed no difference between ΔΕTreg and control skin on P28 (**Figure 2J**). Additionally, Teff numbers decreased in ΔETreg skin on P9, whilst the abundance and proliferation of other T cell subsets were unchanged (**Figure 2L-M**). Nonetheless, we observed an increased IFNγ production by CD8^+^ T cells in the skin on P9 following ETreg depletion (**Supplementary figure 5A-E**), and elevated levels of IFNγ and TNFα in SDLNs (**Supplementary figure 5F-I**). To determine if CD8^+^ T cells play a functional role in ETreg-mediated skin pigmentation, we co-depleted Tregs with CD8^+^ T cells to determine if MeSC function could be rescued in the absence of Tregs. Under these conditions, skin pigmentation was not restored, suggesting that suppression of CD8-mediated inflammation is not a major mechanism by which ETregs promote MeSC function (**Supplementary figure 6A**). Taken together, we are unable to detect any overt T cell-mediated inflammation in ΔETreg mice, either immediately on P9 or in later life on P28, that contributes to the defective skin pigmentation phenotype.

### Melanocytes and PPAR**γ** pathway are under the control of ETregs

We next set out to elucidate the cellular and molecular mechanisms responsible for ETreg-mediated control over MeSC function. We were intrigued by the requirement of Tregs during a very short 3-day window from P6-8, and the lack of fulminant inflammation in ΔETreg mice. Supporting our findings are previous studies showing that adult tissue-resident Tregs facilitate tissue homeostasis largely independently of suppressing conventional inflammatory responses; namely Jagged-1 expressing skin Tregs that promote hair regeneration and Areg-producing Tregs that support lung repair (*10*, *13*). We therefore hypothesized that ETregs regulate either i) the recruitment of specific immune cell subsets we have not assessed in our immune profiling, or ii) specific cellular pathways in non-immune cells, thereby enabling MeSCs to effectively synthesize melanin and maintain postnatal skin homeostasis.

To test this hypothesis, we took a discovery-based approach and reasoned that defining the immediate transcriptomic changes under ETreg control would be key to elucidating the pathways contributing to defective skin pigmentation. We therefore performed bulk RNA-sequencing of whole skin taken from Treg-sufficient controls, ΔETreg, ΔLTreg, and ΔTreg groups 24 hours after the last DT treatment – P9 for ΔETreg or P13 for ΔLTreg and ΔTreg groups (**Figure 3A**). We conducted pathway enrichment analysis using differentially expressed genes (p<0.05) between P9 and P13 skin with ShinyGO (*14*). Interestingly, terms such as melanin biosynthesis, melanin metabolic processes, and melanocyte differentiation showed greater than 15-fold enrichment in P9 controls relative to ΔETreg skin. Additionally, transcriptomic defects in interferon-β responses were evident in the ΔETreg group (**Figure 3B**). In comparison, ΔLTreg skin displayed changes primarily associated with cell development and morphogenesis (**Figure 3C**). These observations suggest that melanocytes are the major cell lineage impacted upon depletion of ETregs, but not LTregs. This notion is further substantiated by the downregulation of key melanocyte transcripts (*Tyr*, *Tyrp1*, *Dct*, *Slc45a2*, *Oca2*, *Slc24a5*) specifically in the ΔETreg and ΔTreg groups, but not in the ΔLTreg group (**Figure 3D**). *In situ* quantification of HF MeSCs using the specific lineage marker CD117 revealed no change in the overall abundance in P9 control or ΔETreg skin, suggesting the transcriptomic changes were not attributed to a quantitative reduction in MeSC numbers (**Figure 3E-F**). Rather, it suggests MeSC function is impacted in ΔETreg animals likely due to downregulation of transcripts associated with melanin synthesis, including *Tyr* and *Oca2*. Overall, we found that melanocytes are key targets of ETregs and that the absence of ETregs lead to disruption of the melanocyte transcriptome. This conclusion is further supported by our histological quantification of MeSC-Treg distance, Teff-Treg and CD8-Treg distances. During early life, Tregs reside in close proximity to MeSCs in the HF region of skin, but not their canonical targets Teffs and CD8^+^ T cells (**Supplementary figure 7A-C**).

**Figure 3.**
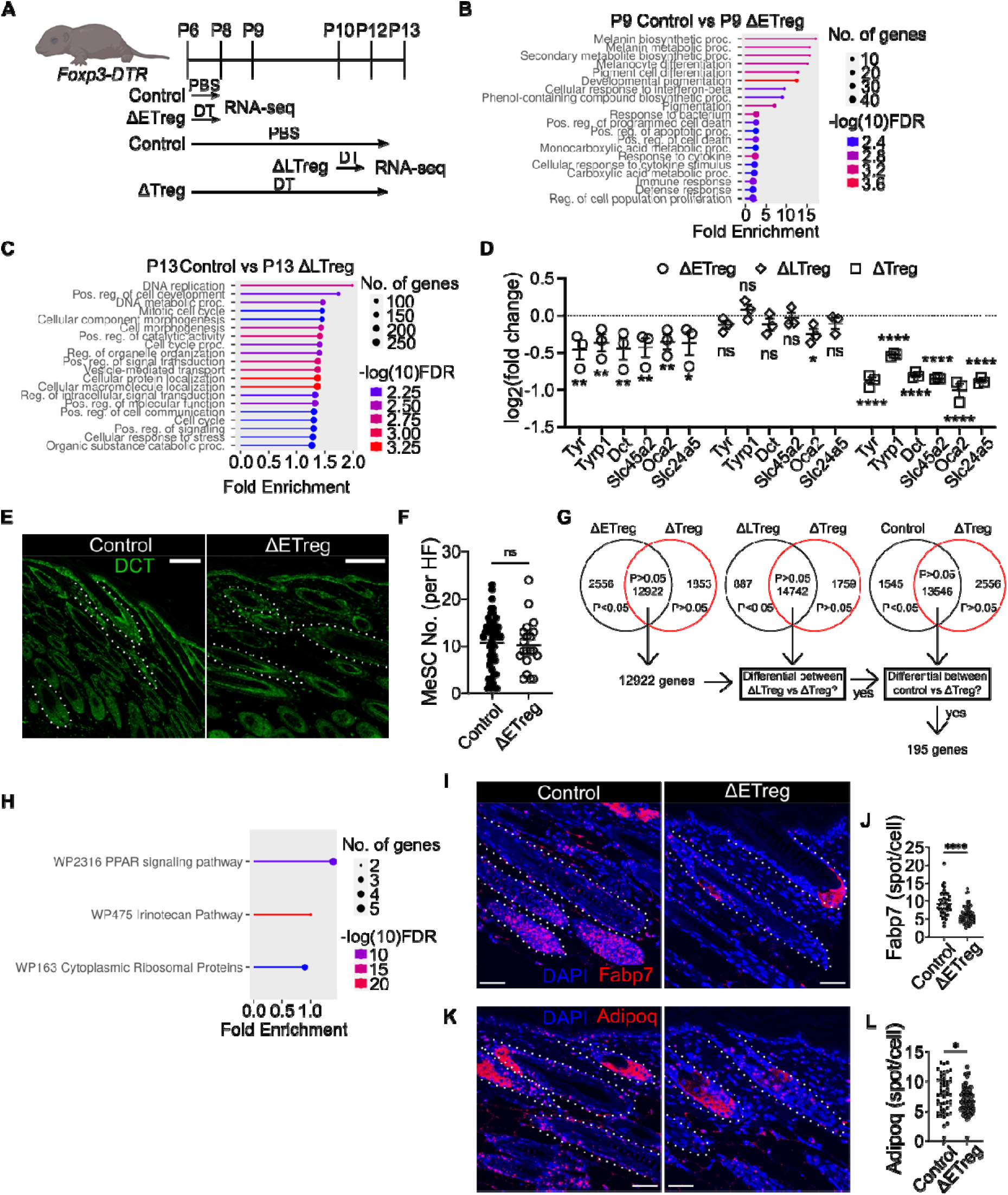
Melanocyte identity and PPARγ activity depend on the presence of early neonatal Tregs. **A)** Schematic outline of experimental timeline. Skin tissues were harvested for bulk RNA-seq on P9 or on P13 depending on treatment group. **B-C)** Enrichment analysis using differentially expressed genes (p<0.05) between age-matched groups. **B)** Control vs ΔETreg skin on P9. **C)** Control vs ΔLTreg on P13. **D)** Normalised expression of melanocyte marker genes. Transcript counts were normalised to age-matched controls (n=2-3 biological replicates). **E)** Immunofluorescence staining of DCT^+^ MeSCs in control and ΔETreg dorsal skin on P9. 100 µm scale bar. **F)** Quantification of MeSC numbers per hair follicle (HF). Unpaired t-test. **G)** Schematic outline of workflow to identify pigmentation-associated genes. 12922 Transcripts that were non-differentially expressed (p>0.05) between ΔETreg and ΔTreg were sequentially filtered. Amongst 12922 transcripts, 195 were also differential (p<0.05) between ΔTreg and ΔLTreg, and ΔTreg and control. **H)** Pathway analysis using pigmentation-associated genes. **I-L)** RNAscope of PPARγ target genes *in situ.* P9 control and ΔETreg skin. 50 µm scale bar. **I)** *Fabp7* and **J**) Quantification of *Fabp7* transcripts in the hair follicle (HF). **K)** *Adipoq* and **L)** Quantification of *Adipoq* transcripts in the HF. **E-L)** Data are representative of 3 independent experiments (n=3-4 biological replicates). Unpaired t-test. ****p<0.0001, **p<0.01, *p<0.05, ns p>0.05. Graphs show mean ± S.E.M.

To identify the signaling pathways responsible for pigmentation downstream of neonatal Tregs, we employed specific inclusion criteria based on the observation that pigmentation fails to develop in the ΔETreg and ΔTreg groups. We selected genes that met the following criteria: 1) Not differentially expressed between the ΔETreg and ΔTreg groups, 2) Differentially expressed between the ΔLTreg and ΔTreg groups, and 3) Differentially expressed between the control and ΔTreg groups (**Figure 3G**). We identified 195 genes that fulfilled these criteria. Pathway enrichment analysis revealed dysregulation of transcripts under the control of the peroxisome proliferator-activated receptor (PPAR) pathway (**Figure 3H**). Amongst the three PPAR isoforms – PPARα, PPARβ, and PPARγ – only PPARγ transitions from high expression to low expression from neonatal to adult skin (*15*). We therefore focused on validation of PPARγ target genes. To validate the immediate decrease in PPARγ target genes, we performed RNAscope analysis of neonatal P9 control and ΔETreg skin to quantify the *in situ* expression of the PPARγ target genes, *Fabp7* and *Adipoq*. The expression of both transcripts specifically in the HFs were significantly attenuated in ΔETreg skin relative to Treg sufficient controls. (**Figure 3I-L**). Overall, our findings convincingly demonstrate that MeSC function in skin is under the tight control of ETregs during neonatal development. We identify a transient window for establishing a Treg-MeSC axis, during which loss of ETregs leads to significant disruption of the PPARγ signalling pathway.

### PPAR**γ** pathway is necessary and sufficient for pigmentation

We next sought to determine if the signalling pathways identified in our transcriptomic analysis are responsible for the pigmentation defect downstream of Tregs. In ΔETreg skin, the two main pathways impacted were associated with Type-I interferon response and PPAR signalling (**Figures 3B**). These candidate pathways have previously been associated with adult melanocyte function *in vitro* (*16*, *17*), but whether they play a role during postnatal skin development *in vivo* has not been assessed. To determine if regulation of the interferon pathway is a major mechanism by which ETregs facilitate MeSC function, we neutralized interferon alpha-receptor (IFNAR) function in ETreg depleted mice using an anti-IFNAR monoclonal antibody. Neutralization of IFNAR was unable to reinstate skin pigmentation in ΔETreg mice, ruling out the interferon-β response as a functional candidate in this process. (**Supplementary figure 6B**).

The next set of experiments addressed if modulation of the PPAR pathway plays a functional role during steady state neonatal skin development. Amongst three isoforms of PPAR (α, β and γ), PPARγ is expressed at higher levels in neonatal skin relative to adult skin (*15*). Therefore, we utilised small molecule modulators targeting the γ isoform. Administration of the highly specific antagonist Τ0070907 to C57BL/6J mice on P6 and P8 significantly impaired skin pigmentation on P28 (**Figure 4B-E).** Defective pigmentation was accompanied by a reduction in the number, but not proliferation, of MeSCs (**Figure 4F-G**). Importantly, these results indicate global PPARγ signalling during neonatal life is required for MeSC-mediated skin pigmentation.

**Figure 4.**
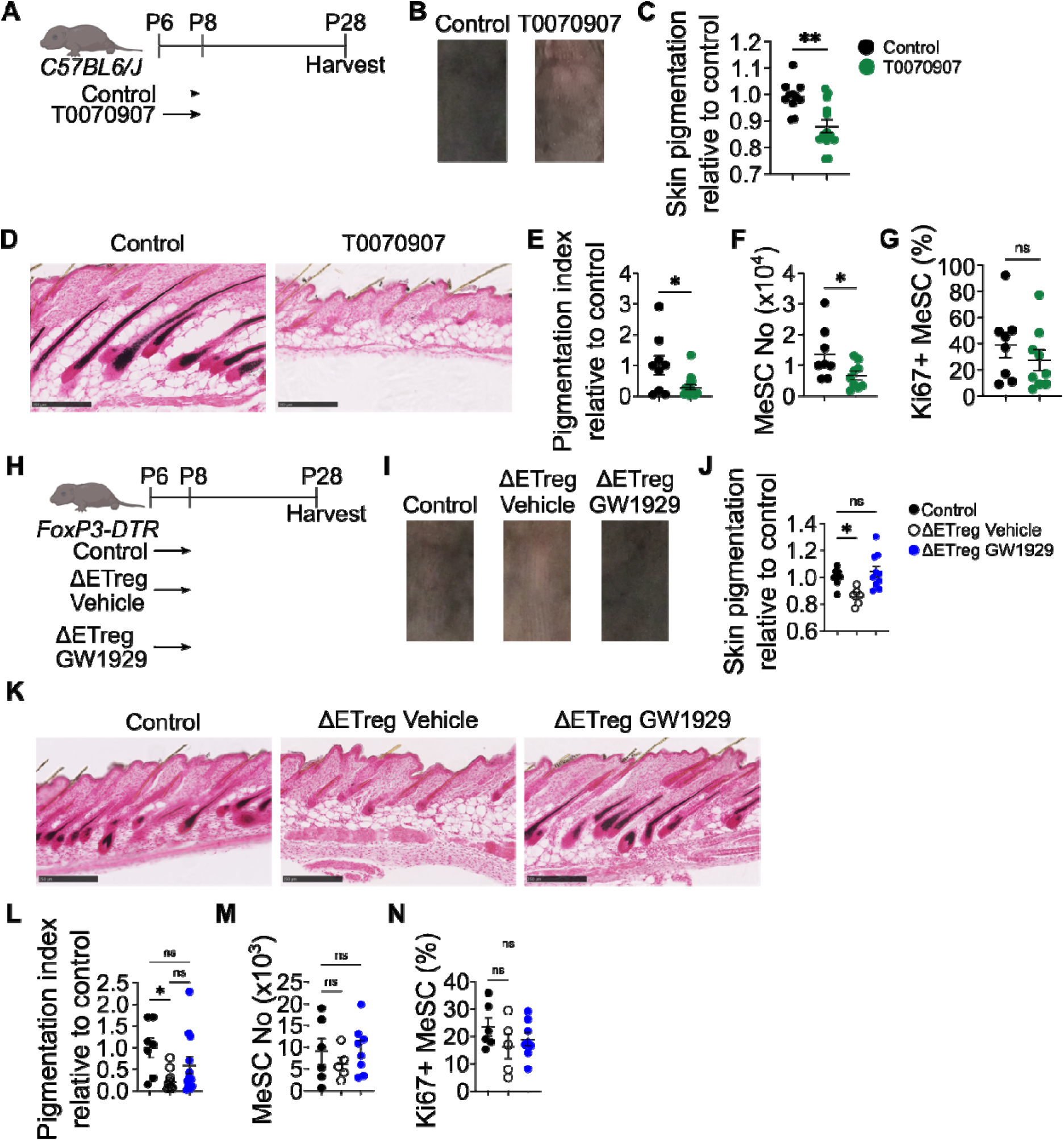
Melanogenic activity requires Treg-PPARγ signalling axis. **A)** Schematic outline of PPARγ antagonization experiment. Control group received DMSO on P6 and P8. T0070907 group received the 2.5 µg/g of PPARγ antagonist, T0070907, on P6 and P8. **B)** Shaved dorsal skin of control and T0070907 mice. **C)** Quantification of skin pigmentation. **D)** Fontana & Masson staining. **E)** Quantification of pigmentation index. **F-G)** Flow cytometric characterisation of MeSCs. **F)** Absolute number. **G)** Percentage of Ki67^+^ MeSCs. **H)** Schematic outline of rescue experiment using PPARγ agonist. Control group received PBS and DMSO on P6 and P8. Τreg-depleted ΔETreg groups received DMSO (vehicle) or GW1929 in addition to DT on P6 and P8. Skin tissues were harvested on P28. **I)** Shaved dorsal skin. **J)** Quantification of skin pigmentation. **K)** Fontana & Masson staining. **L)** Quantification of pigmentation index. Pigmentation was normalised to control. **M-N)** Flow cytometric characterisation of MeSCs. **M)** Absolute number. **N)** Proliferating Ki67^+^ MeSCs. **C, E-G)** Unpaired t-test. **J, L-N)** One way ANOVA against control. Graphs show mean ± S.E.M. Data are pooled from 2 independent experiments (n=4-6 biological replicates). ***p<0.001, **p<0.01, *p<0.05, ns p>0.05.

Next, tο validate whether PPARγ functions downstream of Tregs, we administered a highly specific PPARγ agonist, GW1929, to ΔETreg animals on P6 and P8 (**Figure 4H**). PPARγ agonization restored skin pigmentation in ETreg-depleted mice to baseline levels (**Figure 4I-L**). However, MeSC numbers and proliferation were unaffected by PPARγ agonization (**Figure 4M-N**), suggesting that either qualitative changes in MeSC function, or the presence of an intermediate PPARγ responsive non-immune cell type, may play a role in promoting melanogenesis. Lastly, the observed PPARγ-dependent changes in pigmentation were not associated with significant alterations in skin-resident T cell abundance or proliferative capacity (**Supplementary figure 8A-H**). Therefore, it seems likely that Treg-PPARγ axis functions independently of T cells. Taken together, our findings suggest that a dominant function of ETregs during postnatal development is to regulate the PPARγ signalling axis to establish MeSC-mediated skin pigmentation in later life.

### Epithelial-intrinsic PPAR**γ** signalling activity relies on the presence of ETregs

To begin to understand the diverse cell states in the skin microenvironment, and to further elucidate the cellular mechanisms underlying the ETreg-PPARγ axis, we performed single cell RNA-sequencing (scRNA-seq) of CD45+ and CD45- cells in the presence and absence of Tregs. We reasoned that since Tregs reside proximal to both HFs and the interfollicular epidermis (*10*, *18*, *19*), early compensatory responses of epithelial resident cell types to ETreg depletion likely precede effects on skin pigmentation in later life. Furthermore, because the expansion of activated self-reactive T cells is observed 3-4 days after Treg ablation (*9*) we sought to avoid these confounding variables by analyzing skin-resident cells 1 day after the last DT treatment, on P9. Also, given skin pigmentation requires the presence of P6-8 ETregs, but not P10-12 LTregs, P9 was a rational timepoint to analyze early transcriptional mechanisms that govern pigmentation downstream of ETregs.

Firstly, we quantified CD45^+^ immune cell subsets to assess the accumulation of the major inflammatory lymphoid and myeloid cell lineages in skin. In support of our flow cytometric profiling on P9 (**Supp Figure 5A-E**), pronounced local immune cell activation and inflammation were not observed following ETreg depletion, even though NK cells were moderately increased (**Figure 5A-B**). We also analyzed CD45^-^ immune cells and identified melanocytes (Mel), epithelial cells (Epithelia), hair follicle cells (HF), and other structural cells such as fibroblasts (Fib) and vascular endothelial cells (Vasc). While the overall abundance of these cells was also minimally affected, the proportions of both Mel and Vasc cell populations appear to expand in ΔETreg skin. (**Figure 5C-D**).

**Figure 5.**
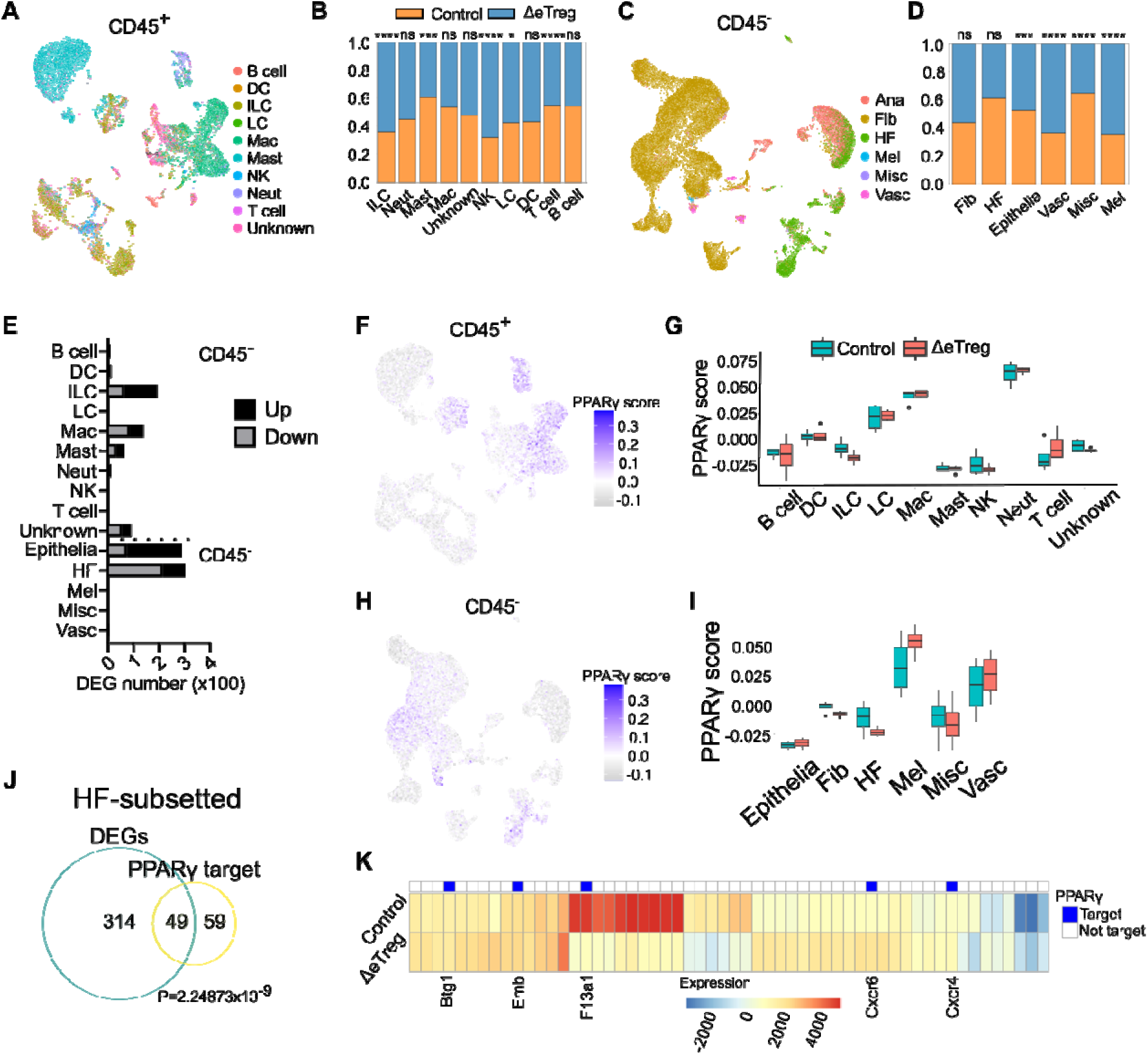
Neonatal Tregs regulate the hair follicle transcriptome and PPARγ activity. **A)** UMAP of CD45^+^ immune cells. B cell. DC, dendritic cell. ILC, innate lymphoid cell. LC, Langerhans cell. Mac, macrophage. Mast, mast cell. NK, natural killer cell. Neut, neutrophil. T cell. Unknown. **B)** Ratio of immune cells in control and ΔETreg skin. **C)** UMAP of CD45^-^ non-immune cells. **D)** Ratio of non-immune cells in control and ΔETreg skin. Epithelia, epithelial cells. Fib, fibroblast. HF, hair follicle. Mel, melanocyte. Misc, miscellaneous. Vasc, vascular cell. **E)** Number of differentially expressed genes (DEG) by cell type. **F-I)** PPARγ activity score. **F-G)** CD45^+^ immune cells. **F)** UMAP representation. **G)** PPARγ score by cell type between control and ΔETreg skin. Η**-**Ι) CD45^-^ non-immune cells. **H)** UMAP representation. **I)** PPARγ score by cell type. **J)** Venn diagram depicting the overlap between the total differentially expressed genes (DEGs) and known PPARγ target genes. P value represents the significance of the overlap as determined by a chi-square test. **K)** Heatmap depicting expression of top differentially expressed genes (LFC>0.5 or LFC<-1).

Next, we performed a pseudo-bulk differential gene expression analysis to ascertain the magnitude of transcriptomic changes induced across all cell populations in ΔETreg and control mice. The most pronounced changes were identified in the HF and Epithelia clusters encompassing all epidermal keratinocytes of skin. When combined, these two cell types account for almost 600 differentially expressed genes (DEGs) (P_adj_<0.05) in the dataset (**Figure 5E and Supplementary Table 2**). Among the hematopoietic CD45^+^ fraction, the innate lymphoid cell (ILC) cluster was most impacted with 200 DEGs imparted by ETreg depletion (**Figure 5E and Supplementary Table 3**). T cell numbers changed minimally, despite being considered the main targets of Treg-mediated suppression. Collectively, these results indicate that the major cellular targets under the control of ETregs in neonatal skin are epithelial and HF cells.

Whole skin bulk RNA-seq and functional *in vivo* skin pigmentation rescue experiments strongly implicate the PPARγ-pigmentation axis as a major mechanism under Treg control (**Figure 3B-L**, **4I-K**). Therefore, we next assessed changes in PPARγ activity across the captured cell types in the presence and absence of ETregs. We selected a set of experimentally validated PPARγ target genes (*20*) and constructed an activity score using the “AddModuleScore” function built into Seurat package. Increasing scores on this scale indicate increased expression of PPARγ target genes. Within the CD45^+^ fraction, neutrophils and macrophages displayed the highest activity levels which is in line with the known role for PPARγ in these cell types (*21*, *22*). However, PPARγ scores in these cell types were largely unaffected by ETreg depletion. Instead, amongst the CD45^-^ cells, the most pronounced downregulation of the PPARγ activity score was observed in HF keratinocytes (**Figure 5I**). This result is in agreement with reduced expression of PPARγ target genes, such as *Fabp7* and *Adipoq* in the HFs (**Figure 3I-L**). This is perhaps an unsurprising result given the HF cluster demonstrated the highest levels of transcriptomic perturbation (**Figure 5E**), and the spatial proximity of HF cells to MeSCs in skin. Further analysis of the HF subset showed that 49 of 363 DEGS (∼14%) between the control and ETreg depleted groups were known transcriptional targets of PPARγ (p= 2.25×10^−9^) (**Figure 5J**). Further visualisation of the top 56 DEGs in HF cells highlight five direct PPARγ targets, two of which are *Cxcr6* and *F13a1* (**Figure 5K**). These transcripts have previously been associated with asymmetric self-renewal capacity of stem cell populations and facilitation of skin wound healing responses, respectively (*23*, *24*). These results indicate that in the absence of ETregs in murine skin, tissue remodelling processes acting via PPARγ signaling is significantly altered in HF cells.

Collectively, scRNA-seq analysis indicates ETreg depletion markedly impacts the abundancies of several non-immune cell subsets in the skin microenvironment while minimally impacting the adaptive lymphoid compartment. Most notably, our results reveal that both the transcriptome and PPARγ activity of HF cells are preferentially modulated by ETregs. As such, neonatal ETregs in skin likely facilitate MeSC function by sustaining PPARγ signalling in HF keratinocytes.

### The PPAR**γ** pathway is implicated in Vitiligo and during human skin development

Numerous disorders of skin pigmentation have been described in humans. The most prominent of which is the autoimmune skin disease, vitiligo, in which depigmented skin results from melanocyte destruction (*25*). Genome wide association studies have highlighted vitiligo susceptibility loci in genes that support Treg function, including IKZF4 and CTLA4 (*26*, *27*), Additionally, Tregs are important for restraining skin depigmentation severity in lesional vitiligo skin, further suggesting a functional role for Tregs in disease pathogenesis. Given these associations, we sought to address whether PPARγ signalling is active in this setting. To test this, we analyzed scRNA-seq data of human skin samples from 5 healthy donors and 10 vitiligo patients (*28*). The final processed datasets yielded 8 main cell types. Consistent with our murine skin single cell data, we identified two major clusters of keratinocytes in healthy and vitiligo skin; ‘HF’ characterized by high expression of hair follicle--associated markers *KRT14* and *KRT15*, and ‘Epithelia’ that expressed the interfollicular epidermal transcript *KRT5* (**Figure 6D**). We next applied the PPARγ scoring module to assess the level of activity across the captured cell populations (*20*). These analyses revealed that only the HF cell type from vitiligo skin exhibited a significant downregulation in PPARγ pathway activation, relative to heathy controls. (**Figure 6E**). This result aligns closely to our murine single cell data where dysregulation of PPARγ signalling was identified in HF cells upon loss of ETregs in neonatal skin (**Figure 5I**).

**Figure 6.**
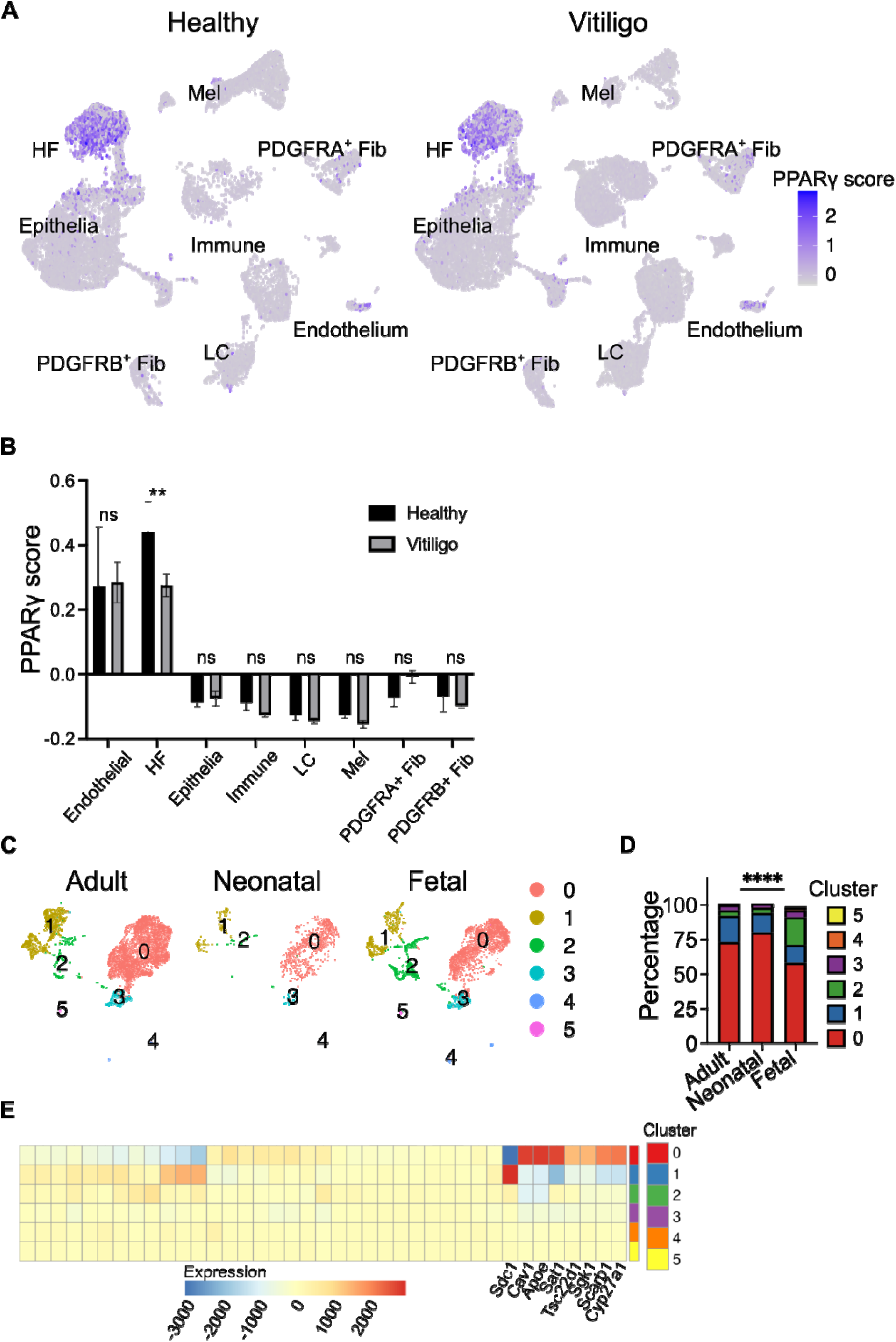
PPARγ target genes are differentially regulated between human developmental stages and diseased states. **A-B)** scRNA-seq analysis of skin from healthy donors and vitiligo patients. **A)** UMAP representation of PPARγ target gene expression score. Endothelial, endothelial cells. HF, hair follicle. IFE, interfollicular epidermis. Immune, immune cells. LC, Langerhans cells. Mel, melanocytes. PDGFRA+ Fib, PDGFRA+ fibroblasts. PDGFR+ Fib, PDGFRB+ fibroblasts. **B)** Quantification of PPARγ score. Fisher’s LSD test. **C-E)** scRNA-seq analysis of fetal, neonatal, and adult melanocytes. **C)** UMAP representation of clusters. **D)** Percentage of cells belonging to individual clusters. Chi-squared test. **E)** Heatmap of PPARγ target gene expression. ****p<0.0001, **p<0.01, ns p>0.05.

We next set out to address if the PPARγ pathway is involved in human neonatal melanocyte development. *In utero*, Tregs seed the skin early as gestational week (GW) 18 (*15*, *29*). This timeframe aligns closely with murine skin development during the first week of life when ETregs begin to accumulate (**Figure 1**). Importantly, this period of Treg infiltration coincides with human HF development, which involves the migration of melanoblasts to the HF to complete the melanocyte maturation process (*30*). Given these associations, we analyzed scRNA-Seq data of melanocytes isolated from fetal (9.5-18 GW), neonatal (0 years), and adult (24-81 years) tissue (*28*, *31*). We identified 6 distinct clusters across the dataset. Chi-squared analysis revealed a marked enrichment of adult and neonatal melanocytes for cluster 0, while fetal melanocytes were enriched for cluster 2 (**Figure 6A-B**). This observation suggests that melanocyte maturation involves the expansion of cluster 0. Indeed, cluster 0 expresses high levels of mature melanocyte markers such as *DCT, PMEL, TYR*. Cluster 2 expresses high level of proliferation-associated transcripts such as *MKI67, TOP2A and CCNB1,* therefore representing actively dividing melanocytes (**Supplementary table 4**). Further analysis identified elevated expression of multiple PPARγ target genes in cluster 0 (**Figure 6C**) (*32*), including *CAV1* (*33*), which regulates melanogenesis and *SCARB1*, which is involved in fatty acid uptake (*34*). Overall, an increased PPARγ activity is associated with melanocyte maturation. The transition frow low PPARγ activity to high PPARγ activity temporally coincides with Treg seeding. As such, our findings suggest the possibility that human Tregs may modulate PPARγ activity during human skin development.

Collectively, our analysis of human transcriptomic data suggests that the PPARγ pathway is active during melanocyte development and is preferentially disrupted in HF cells from the pigment-deficient disease state Vitiligo.

## DISCUSSION

Conventionally, Tregs are defined by their robust capacity to dampen tissue inflammatory responses. More recently, there is an increasing realization of the important non-conventional role of adult Tregs in supporting normal tissue maintenance and repair that is either independent of, or mutually codependent with, control of local inflammation (*5*, *10*, *11*, *13*, *18*, *35*, *36*). In neonates, Tregs are instrumental in establishing mechanisms of peripheral tolerance to both tissue-specific and commensal bacterial antigens (*6*, *7*), as well as regulation of the skin stromal cell niche during the second week of life (*5*). Furthermore, adult Tregs support the bone marrow and skin stem cell (SC) niche (*10*, *37*, *38*), but whether a Treg-SC axis exists in neonatal tissue has not been established.

Here, we show that early skin seeding Tregs (ETregs) in the first week of life play a crucial role in establishing tissue homeostatic processes that manifest in later life. We observed fluctuations of the critical Treg effector molecules CD25 and CTLA4 between postnatal day 6 (P6), P9, and P12 skin, suggesting a temporal dynamic in Treg function during early neonatal stages. Indeed, transient depletion of P6-P8 ETregs, but not P10-P12 late skin seeding Tregs (LTregs), led to impaired melanocyte stem cell (MeSC) mediated skin pigmentation on P28. Under these conditions, whole skin transcriptomic analysis identified melanocyte dysfunction and modulation of the PPARγ pathway as under ETreg control. These changes occurred immediately, within 24 hours, following ETreg depletion before any signs of a fulminant skin inflammatory response. Restoration of neonatal PPARγ signalling re-instated skin pigmentation, suggesting that this pathway plays a major role downstream of ETregs. By contrast, LTreg depletion leads to transcriptomic changes in genes associated with tissue morphogenesis, suggesting that LTregs may regulate the structural integrity of the skin (**Figure 3C**). In line with this observation, depletion of late-seeding Tregs on P8 and P15 leads to the absence of dermal white adipose tissue in later life (*5*). We observe similar loss of adipose tissues upon a 4-dose depletion of neonatal Tregs on P6, P8, P10 and P12 (**Supplementary Figure 3C**). Together, these observations strongly support temporal dynamic in Treg function. Our flow cytometric profiling of Treg activation markers, such as CD25 and CTLA4, suggest that an intrinsic functional difference between early and late neonatal Tregs are responsible for distinct phenotypes produced by temporal variation in depletion regimen. Another plausible explanation for the phenotypic differences is that the rapidly developing postnatal skin is only dependent on ETregs for melanogenic activity.

In addition to its immunomodulatory functions, the PPARγ pathway plays a key role in the accumulation and adaptation of Tregs within adipose tissue microenvironments, such as the dermal white adipose underlying the dermis (*39*). However, skin T cell numbers were unaffected by transient antagonization of the PPARγ pathway in neonatal mice (**Supplementary figure 8B**). This suggests that the Treg-melanocyte axis may operate independently of skin-resident adaptive immune cells. Moreover, skin pigmentation was not restored in ΔETreg mice upon depletion of CD8^+^ T cells or blockade of the interferon-β response (**Supplementary figure 6A-B**). Whether the promotion of melanocyte function is mediated by a subset of skin-resident ETregs or the bulk ETreg population, and whether Treg expression of a tissue specific factor(s) is a major mechanism that ETregs utilize remains to be determined. Together, these observations support the idea that suppression of inflammation is not the major mechanism by which ETregs promote melanocyte mediated skin pigmentation.

Finally, our whole tissue single cell analysis suggests that Tregs preferentially regulate PPARγ activity within the HF epithelium (**Figure 3I-L**, **5I**). Globally, the transcriptome of HFs was impacted most significantly relative to other cell types in skin immediately upon depletion of ETregs (**Figure 5E**). Similar findings were reported in a recent study assessing the transient loss of Tregs in the contexts of lung cancer and injury induced inflammation. Within 2 days of Treg ablation there were pronounced transcriptional responses in stromal cells, but not adaptive immune cells (*40*). The authors demonstrate that this Treg ‘connectivity’ to non-haematopoietic cell types in lung is facilitated by *in situ* proximity to these first responding cell types. Our finding that ETregs reside in markedly closer proximity to MeSCs within the HF regions than CD8^+^ T cells or CD4^+^ Teffs supports the notion that Tregs are highly connected to HFs in skin (**Supplementary figure 7A-C**) (*18*). We also demonstrate the relevance of this connection in the human skin pigmentation disorder, vitiligo, where Tregs have been implicated in restraining clinical severity. Our analysis uncovers preferential disruption of PPARγ signaling within the HF epithelium in diseased skin relative to healthy controls (**Figure 6B**). Furthermore, in developing human skin we show that melanocytes are enriched for this pathway at gestational week 9.5-18 (**Figure 6C-E**). Interestingly, this time-frame coincides almost precisely with Treg seeding of fetal human skin (*41*). Together, these observations suggest that Tregs during very early life are functionally and spatially poised to modulate PPARγ responses in the HF epithelium and that modulation of this pathway is also impacted in human diseased skin where Tregs play a functional role.

A major limitation of our study is the use of PPARγ modulators, instead of transgenic mice. Melanocyte abundance is reduced in wild-type mice following neonatal administration of the antagonist T0070907, but not in ΔETreg mice (**Figure 2F**, **4F**). Therefore, it is possible that distinct mechanisms lead to pigmentation defects in these two contexts. This discrepancy may be explained by our observation that melanocytes are most dependent on PPARγ signalling activity in developing mouse skin, and therefore most vulnerable to PPARγ antagonists (**Figure 5G**).

Overall, our study provides new insights into the function of the foremost skin-seeding Tregs during neonatal life. Immune profiling during the first 12 days indicates a temporal dynamic in Treg function, whereby early skin seeding Tregs, but not later seeding Tregs, potentiate PPARγ signalling to facilitate melanocyte behaviour. Pertinently, GWAS studies have implicated Treg malfunction in the human melanocyte-associated disease, vitiligo (*42*). Future studies assessing human neonatal skin Treg frequency, phenotype and perhaps most critically, comprehensive single cell-to-cell spatial co-localization with stem cell populations in the HF epithelia may uncover previously unrecognized Treg modalities during early life that may contribute to disease pathogenesis in later life.

## MATERIALS AND METHODS

### Mouse studies

All animal experiments were subject to local ethical approval and performed under the terms of a UK government Home Office licence (PP6051479). All mice were outbred on a C57BL/6J background. Both male and female mice were used. B6.129(Cg)-Foxp3^tm3(DTR/GFP)^Ayr/J (Foxp3-DTR) mice were purchased from Jackson laboratories. For Treg depletion, *Foxp3-DTR* mice intraperitoneally received 30 ng/g of DT on P6 and P8. PPARγ antagonization was performed by intraperitoneal administration of 2.5 µg/g of T0070907 on P6 and P8. Rescue studies were performed by intraperitoneal administration of 30 ng/g of DT and 20 µg/g of GW1929 on P6 and P8. CD8^+^ T cells were depleted by intraperitoneal administration of 100 µg of α-CD8 depleting antibody (BioXcell) on P6, P8, P15 and on P22. Interferon-β responses were blocked by intraperitoneal administration of α-Ifnar blocking antibody (Biolegend) on P6 and P8. Pre-weanlings were sacrificed by overdose of pentobarbital. Post-weanlings were sacrificed by CO2 asphyxiation followed by severing a major blood vessel. All efforts were made to minimise suffering for mice.

### Tissue digestion

Single-cell suspensions of full-thickness skin for flow cytometry was performed as previously described (*10*). Isolation of cells from axillary, brachial and inguinal lymph nodes for flow cytometry was performed by mashing tissue over 70 μm sterile filters. To prepare dorsal skin cell suspension, shaved mouse skin was de-fatted, minced finely with scissors, and re-suspended in a 3 ml of C10 (RPMI-1640 with L-glutamine with 10% heat-inactivated FBS, 1% penicillin-streptomycin, 1 mM Sodium pyruvate, 1% Hepes, 1x Non-essential amino acid and 60 μM β-mercaptoethanol) supplemented with 2mg/ml collagenase XI, 0.5mg/ml hyaluronidase and 0.1mg/ml DNase in a 50 ml conical. The mixture was digested in a shaking incubator at 37°C at 255 rpm for 45 mins. After a vigorous shaking, the digested mixture was passed through a sterile 100 μm filter fitted onto a 50 ml conical. After pelleting, the filtrate was then filtered once more through a 40 μm strainer fitted onto a new 50 ml conical. Finally, the cells were pelleted once more and re-suspended in 1 ml of C10. Epidermal cell suspensions were prepared by scraping the dermis away with forceps and incubating the layer of epidermis on 3 mL of 0.5% Trypsin-EDTA (ThermoFisher) at 37°C for 1 hour. Epidermal cells were isolated by scraping the trypsinised epidermis on a petri dish containing 4 ml of C10 media. The mixture of cell suspension and scraped skin was filtered through a 70 μm strainer fitted onto a 50 ml conical. Finally, the cells were pelleted and re-suspended in C10. After cell count using nucleocounter (Chememoetec), cells were plated on nunc round bottom 96-well plates (Thermofisher) for staining.

### Flow cytometry

Following isolation from the tissue, cells were labeled stained in PBS for 20 mins at 4°C with a live dead marker (Zombie UV^TM^, Biolegend). Surface staining was performed in brilliant stain buffer (BD Biosciences) for 20 mins at 4°C. For intracellular staining, cells were fixed and permeabilized using reagents and protocol from the Foxp3 staining buffer kit (eBioscience). Fluorophore-conjugated antibodies specific for mouse cell surface antigens and intracellular transcription factors were purchased from eBioscience, BD Biosciences or Biolegend as detailed in the Supplementary Table 1. All samples were run on Fortessa LSRII (BD Biosciences) at the KCL BRC Flow Cytometry Core. Experiments were standardised using SPHERO Rainbow calibration particle, 8 peaks (BD Biosciences, 559123). For compensation, UltraComp eBeads (Thermo Fisher, 01-2222-42) were stained for each surface and intracellular antibody following the same procedure as cell staining. ArC Amine Reactive Compensation Bead Kit (Thermo Fisher, A10346) were used for Zombie UV^TM^ stain. All gating and data analysis were performed using FlowJo v10, while statistics were calculated using GraphPad Prism 9. Strict dead cell and doublet cell exclusion criteria were included for all immune cell analysis, followed by pre-gating for all hematopoietic cells as CD45^+^. All immune cells were pre-gated as Zombie UV^-^ CD45^+^ cells. CD3^-^ lymphoid/myeloid cells were gated TCR^γδ–^CD3^−^ double negative cells. Other lymphoids were gated as TCRγδ^+^CD3^+^ dermal γδ T cells (dGDTCs), TCRγδ^hi^CD3^hi^ dendritic epidermal T cells (DETCs), ΤCDγδ^-^CD3^+^CD8^+^ T cells (CD8), ΤCRγδ^-^ CD3^+^CD4^+^Foxp3^−^ T effector cells (Teff), and ΤCRγδ^-^CD3^+^CD4^+^Foxp3^+^ regulatory T cells (Tregs).

### T cell stimulation

PMA/Ionomycin cocktail (Tonbo Biosciences) was diluted to 1x concentration in C10. Heat-inactivated FBS must be used to make C10 for cell stimulation. 6 to 8 million live cells were re-suspended in 200 μl of 1x PMA/Ionomycin cocktail and plated on a round-bottom 96-well plate (Thermofisher Scientific). Re-suspended cells were incubated at 37°C for 4 hours. Stimulated cells were centrifuged at 1800 rpm for 4 minutes at 4°C. Cell pellets were washed by topping up with 200 μl of FACS buffer and centrifugation at 1800 rpm for 4 minutes at 4°C. Washed cells were stained as follows: live/dead staining for 20 minutes on ice; surface staining overnight at 4°C; and intracellular staining overnight at 4°C.

### Histology and microscopy

Loose fatty tissues were removed from shaved dorsal skin with a pair of forceps. The skin was fixed in 10% neutral buffered formalin (Fisher Scientific) overnight at 4°C. On the following day, fixed tissues were washed twice in PBS for 5 mins each on a rocking platform. Following wash, skin tissues were either stored in 70% EtOH at 4°C to later be embedded in paraffin. Fontana & Masson staining was performed as per manufacturer’s protocol (Abcam). F&M slides were imaged using a Nanozoomer (Hamamatsu photonics) with a ×40 objective.

### Immunofluorescence and tissue microarray

Immunofluorescence labelling was performed using tissue microarrays generated using a manual tissue arrayer (Beechers Instruments). Individual tissue cores (4 mm diameter) were extracted from FFPE skin tissue blocks and moved to a recipient block using a Beechers MTA1 manual tissue arrayer (Beechers Instruments). Multiplex staining was performed using the following antibodies anti-FoxP3 (14-5773-82, Thermofisher), anti-DCT (ab74073, Abcam), and anti-CD117 (AF1356, RnD systems). Immunohistochemical experiments were conducted using Bond Max automated staining system (Leica, UK). Each antibody was optimized for pH and concentration dependence, antigen retrieval and temperature parameters. RNAscope was performed as per manufacturer’s protocol. Confocal microscopy was performed with a Leica SP8 confocal microscope using a ×20 objective.

### *In situ* transcript quantification

RNA-scope slides were analyzed using QuPath (*43*). Cells were detected using the DAPI channel. Transcript counts per cell was quantified using built-in subcellular spot detection function.

### Pigmentation index

Pigmentation index was calculated from F&M images as follows. Images were converted to 16-bit using Fiji. Otsu method thresholding was performed to filter for black pixels corresponding to melanin. Area of melanin was measured and normalised to the length of the skin.

### Whole skin RNA-seq

4 mm biopsy of skin samples were stored in RNAlater (Sigma) at -20°C. Skin samples were homogenised with Kinematica^TM^ Polytron^TM^ handheld PT1200E homogeniser (Fisher Scientific) using a 1200E probe (Fisher Scientific). The RNA was extracted using the mirVana^TM^ miRNA Isolation kit (Invitrogen) by following the manufacturer’s protocol. Prior to start of digestion, and between samples, the probe was sequentially washed in 1% SDS, 100% ethanol and dH2O to avoid cross-contamination. Samples were maintained on ice at all times. Extracted RNA was stored at -20°C. Concentration of the RNA was determined using Qubit^TM^ Broad Range assay kit (Invitrogen). Quality of the RNA was checked using Bioanalyzer (Agilent Technologies) at the King’s Genomics Centre. Only samples with RNA integrity score greater than 7.0 were taken further for downstream applications. 2 μg of total RNA per samples were used for RNA-seq. 150 bp paired-end Sequencing was performed on the NovaSeq platform (Illumina) at Genewiz. Sequenced reads were trimmed using Trimmomatic v.0.36. Trimmed reads were mapped to the Mus musculus GRCm38 reference genome using the STAR aligner v.2.5.2b to produce BAM files. Unique gene counts were calculated using featureCounts function from the Subread package v.1.5.2. Only reads that fell within exon regions were counted to generate the gene hit counts table. Differential expression analysis was performed using DESeq2. Wald test was used to generate p-values and log2 fold changes.

### Pathway analysis

An enrichment analysis was performed to extract biological information from the list of pigmentation-associated genes. The enrichment analysis was performed using a web-based platform, ShinyGO (136). A false discovery rate cut-off of 0.3 was used for the list of pigmentation-associated genes. All other pathway analysis was performed with a false discovery rate cut-off of 0.05.

### Mouse skin scRNA-seq

Skin CD45^+^ and CD45^-^ cells were FACS-sorted into RPMI-1640 supplemented with 40% heat-inactivated FBS and 40 U/ml of RiboLock RNase inhibitor (Thermofisher). The cells numbers were counted and mixed at 2:1 ratio (CD45^+^:CD45^-^). The mixed cells were used for cDNA library generation using the 10X Genomics Chromium Sinlge cell 3’ kit by the Genomics Centre at the Blizard Institute. Prepared libraries were sequenced using the NovaSeq 6000 S4. Raw sequencing data were processed using the 10X Genomics *CellRanger* package.

### Human skin scRNA-seq

Human neonatal and adult melanocyte scRNA-seq data is available in the Gene Expression Omnibus (GEO) database repository under accession number GSE151091. Human healthy and vitiligo skin scRNA-seq data is available in the Genome Sequence Archive (GSA) with accession number PRJCA006797. Analysis was performed using Seurat package in the R programming environment.

### Statistical analysis

Statistical analysis was performed using GraphPad Prism 8.0 (GraphPad Inc.). Unpaired t-test, paired t-test or one way ANOVA was performed as indicated in figure legends to calculate p values. Statistical significance was inferred if p value was less than 0.05 (*), 0.01 (**), 0.001 (***), or 0.0001 (****). P values greater than 0.05 was identified as not statistically significant.

## Supplementary Materials

Supplementary figure 1. Flow cytometric gating strategy for analysis.

Supplementary figure 2. Flow cytometric characterisation of T cell numbers and proliferation.

Supplementary figure 3. Four-dose DT regimen causes weight loss, inflammation, and pigment defect.

Supplementary figure 4. Transient DT injection is sufficient for skin Treg depletion.

Supplementary figure 5. Early Treg depletion causes inflammatory response in the skin and SDLN.

Supplementary figure 6. Modulation of skin pigmentation.

Supplementary figure 7. Skin Tregs reside proximally to melanocyte stem cells, and not T cells.

Supplementary figure 8. Modulation of neonatal PPARγ signalling does not affect skin-resident immune cell numbers or proliferation.

Supplementary table 1. List of antibodies for flow cytometry.

Supplementary table 2. Differentially expressed gene list from single CD45^-^ cell RNA-seq analysis.

Supplementary table 3. Differentially expressed gene list from single CD45^+^ cell RNA-seq analysis.

Supplementary table 4. Cluster markers from developing human melanocyte single cell RNA- seq.

## Acknowledgments

We gratefully acknowledge the Advanced Cytometry Platform (Flow Core), Research and Development Department at Guy’s and St Thomas’ NHS Foundation Trust, the imaging facility staff at University College London, Barts Cancer Institute Flow Cytometry Facility and the Genomics Centre at the Blizard Institute at Queen Mary University of London. We also thank Hee-Yeon Jeon and Jinwook Choi for helpful discussions.

## Author contributions

Conceptualization: NA

Methodology: IC, BZ, NA

Investigation: IC, JX, PL, HA, SA

Visualization: IC, PT, HJ, JC

Funding acquisition: NA

Project administration: NA

Supervision: NA

Writing – original draft: IC, NA

Writing – review & editing: IC, NA

## References

1. I. Cho, P. P. Lui, N. Ali, Treg regulation of the epithelial stem cell lineage. J. Immunol. Regen. Med. (2020), doi:10.1016/j.regen.2020.100028.

2. P. P. W. Lui, I. Cho, N. Ali, Tissue regulatory T cells. Immunology (2020),, doi:10.1111/imm.13208.

3. M. Panduro, C. Benoist, D. Mathis, Tissue Tregs. Annu. Rev. Immunol. (2016), doi:10.1146/annurev-immunol-032712-095948.

4. T. C. Scharschmidt, K. S. Vasquez, M. L. Pauli, E. G. Leitner, K. Chu, H. A. Truong, M. M. Lowe, R. Sanchez Rodriguez, N. Ali, Z. G. Laszik, J. L. Sonnenburg, S. E. Millar, M. D. Rosenblum, Commensal Microbes and Hair Follicle Morphogenesis Coordinately Drive Treg Migration into Neonatal Skin. Cell Host Microbe (2017), doi:10.1016/j.chom.2017.03.001.

5. I. C. Boothby, M. J. Kinet, D. P. Boda, E. Y. Kwan, S. Clancy, J. N. Cohen, I. Habrylo, M. M. Lowe, M. Pauli, A. E. Yates, J. D. Chan, H. W. Harris, I. M. Neuhaus, T. H. McCalmont, A. B. Molofsky, M. D. Rosenblum, Early-life inflammation primes a T helper 2 cell–fibroblast niche in skin. Nature (2021), doi:10.1038/s41586-021-04044-7.

6. T. C. Scharschmidt, K. S. Vasquez, H. A. Truong, S. V. Gearty, M. L. Pauli, A. Nosbaum, I. K. Gratz, M. Otto, J. J. Moon, J. Liese, A. K. Abbas, M. A. Fischbach, M. D. Rosenblum, A Wave of Regulatory T Cells into Neonatal Skin Mediates Tolerance to Commensal Microbes. Immunity (2015), doi:10.1016/j.immuni.2015.10.016.

7. S. Yang, N. Fujikado, D. Kolodin, C. Benoist, D. Mathis, Regulatory T cells generated early in life play a distinct role in maintaining self-tolerance. Science (80-.). (2015), doi:10.1126/science.aaa7017.

8. P. Sivasami, C. Elkins, P. P. Diaz-Saldana, K. Goss, A. Peng, M. Hamersky IV, J. Bae, M. Xu, B. P. Pollack, E. M. Horwitz, C. D. Scharer, L. Seldin, C. Li, Obesity-induced dysregulation of skin-resident PPARγ^+^ Treg cells promotes IL-17A-mediated psoriatic inflammation. Immunity. 56, 1844–1861.e6 (2023).

9. J. M. Kim, J. P. Rasmussen, A. Y. Rudensky, Regulatory T cells prevent catastrophic autoimmunity throughout the lifespan of mice. Nat. Immunol. 8, 191–197 (2007).

10. N. Ali, B. Zirak, R. S. Rodriguez, M. L. Pauli, H. A. Truong, K. Lai, R. Ahn, K. Corbin, M. M. Lowe, T. C. Scharschmidt, K. Taravati, M. R. Tan, R. R. Ricardo-Gonzalez, A. Nosbaum, M. Bertolini, W. Liao, F. O. Nestle, R. Paus, G. Cotsarelis, A. K. Abbas, M. D. Rosenblum, Regulatory T Cells in Skin Facilitate Epithelial Stem Cell Differentiation. Cell (2017), doi:10.1016/j.cell.2017.05.002.

11. A. N. Mathur, B. Zirak, I. C. Boothby, M. Tan, J. N. Cohen, T. M. Mauro, P. Mehta, M. M. Lowe, A. K. Abbas, N. Ali, M. D. Rosenblum, Treg-Cell Control of a CXCL5-IL-17 Inflammatory Axis Promotes Hair-Follicle-Stem-Cell Differentiation During Skin-Barrier Repair. Immunity (2019), doi:10.1016/j.immuni.2019.02.013.

12. J. E. Harris, T. H. Harris, W. Weninger, E. J. Wherry, C. A. Hunter, L. A. Turka, A mouse model of vitiligo with focused epidermal depigmentation requires IFN-γ for autoreactive CD8+ T-cell accumulation in the skin. J. Invest. Dermatol. (2012), doi:10.1038/jid.2011.463.

13. N. Arpaia, J. A. Green, B. Moltedo, A. Arvey, S. Hemmers, S. Yuan, P. M. Treuting, A. Y. Rudensky, A Distinct Function of Regulatory T Cells in Tissue Protection. Cell (2015), doi:10.1016/j.cell.2015.08.021.

14. S. X. Ge, D. Jung, D. Jung, R. Yao, ShinyGO: A graphical gene-set enrichment tool for animals and plants. Bioinformatics (2020), doi:10.1093/bioinformatics/btz931.

15. L. Michalik, B. Desvergne, N. S. Tan, S. Basu-Modak, P. Escher, J. Rieusset, J. M. Peters, G. Kaya, F. J. Gonzalez, J. Zakany, D. Metzger, P. Chambon, D. Duboule, W. Wahli, Impaired skin wound healing in peroxisome proliferator-activated receptor (PPAR)α and PPARβ mutant mice. J. Cell Biol. (2001), doi:10.1083/jcb.200011148.

16. K. Krasagakis, C. Garbe, S. Krüger, C. E. Orfanos, Effects of interferons on cultured human melanocytes in vitro: Interferon-beta but not -alpha or -gamma inhibit proliferation and all interferons significantly modulate the cell phenotype. J. Invest. Dermatol. (1991), doi:10.1111/1523-1747.ep12480767.

17. J. S. Lee, Y. M. Choi, H. Y. Kang, PPAR-gamma agonist, ciglitazone, increases pigmentation and migration of human melanocytes. Exp. Dermatol. (2007), doi:10.1111/j.1600-0625.2006.00521.x.

18. J. M. Moreau, M. O. Dhariwala, V. Gouirand, D. P. Boda, I. C. Boothby, M. M. Lowe, J. N. Cohen, C. E. Macon, J. M. Leech, L. A. Kalekar, T. C. Scharschmidt, M. D. Rosenblum, Regulatory T cells promote innate inflammation after skin barrier breach via TGF-β activation. Sci. Immunol. (2021), doi:10.1126/sciimmunol.abg2329.

19. I. K. Gratz, H.-A. Truong, S. H.-Y. Yang, M. M. Maurano, K. Lee, A. K. Abbas, M. D. Rosenblum, Cutting Edge: Memory Regulatory T Cells Require IL-7 and Not IL-2 for Their Maintenance in Peripheral Tissues. J. Immunol. (2013), doi:10.4049/jimmunol.1300212.

20. L. Fang, M. Zhang, Y. Li, Y. Liu, Q. Cui, N. Wang, PPARgene: A Database of Experimentally Verified and Computationally Predicted PPAR Target Genes. PPAR Res. 2016, 6042162 (2016).

21. P. Tontonoz, L. Nagy, J. G. A. Alvarez, V. A. Thomazy, R. M. Evans, PPARγ Promotes Monocyte/Macrophage Differentiation and Uptake of Oxidized LDL. Cell. 93, 241–252 (1998).

22. T. J. Standiford, V. C. Keshamouni, R. C. Reddy, “Peroxisome proliferator-activated receptor-γ as a regulator of lung inflammation and repair” in Proceedings of the American Thoracic Society (2005).

23. M. Noh, J. L. Smith, Y. H. Huh, J. L. Sherley, A resource for discovering specific and universal biomarkers for distributed stem cells. PLoS One (2011), doi:10.1371/journal.pone.0022077.

24. G. Theocharidis, D. Baltzis, M. Roustit, A. Tellechea, S. Dangwal, R. S. Khetani, B. Shu, W. Zhao, J. Fu, S. Bhasin, A. Kafanas, D. Hui, S. H. Sui, N. A. Patsopoulos, M. Bhasin, A. Veves, Integrated Skin Transcriptomics and Serum Multiplex Assays Reveal Novel Mechanisms of Wound Healing in Diabetic Foot Ulcers. Diabetes. 69, 2157–2169 (2020).

25. M. L. Frisoli, J. E. Harris, Vitiligo: Mechanistic insights lead to novel treatments. J. Allergy Clin. Immunol. (2017), doi:10.1016/j.jaci.2017.07.011.

26. N. Ohkura, Y. Yasumizu, Y. Kitagawa, A. Tanaka, Y. Nakamura, D. Motooka, S. Nakamura, Y. Okada, S. Sakaguchi, Regulatory T Cell-Specific Epigenomic Region Variants Are a Key Determinant of Susceptibility to Common Autoimmune Diseases. Immunity. 52 (2020), doi:10.1016/J.IMMUNI.2020.04.006.

27. Y. Jin, G. Andersen, D. Yorgov, T. M. Ferrara, S. Ben, K. M. Brownson, P. J. Holland, S. A. Birlea, J. Siebert, A. Hartmann, A. Lienert, N. van Geel, J. Lambert, R. M. Luiten, A. Wolkerstorfer, J. P. Wietze van der Veen, D. C. Bennett, A. Taïeb, K. Ezzedine, E. H. Kemp, D. J. Gawkrodger, A. P. Weetman, S. Kõks, E. Prans, K. Kingo, M. Karelson, M. R. Wallace, W. T. McCormack, A. Overbeck, S. Moretti, R. Colucci, M. Picardo, N. B. Silverberg, M. Olsson, Y. Valle, I. Korobko, M. Böhm, H. W. Lim, I. Hamzavi, L. Zhou, Q.-S. Mi, P. R. Fain, S. A. Santorico, R. A. Spritz, Genome-wide association studies of autoimmune vitiligo identify 23 new risk loci and highlight key pathways and regulatory variants. Nat. Genet. 48, 1418–1424 (2016).

28. Z. Xu, D. Chen, Y. Hu, K. Jiang, H. Huang, Y. Du, W. Wu, J. Wang, J. Sui, W. Wang, L. Zhang, S. Li, C. Li, Y. Yang, J. Chang, T. Chen, Anatomically distinct fibroblast subsets determine skin autoimmune patterns. Nature (2022), doi:10.1038/s41586-021-04221-8.

29. M. O. Dhariwala, D. Karthikeyan, K. S. Vasquez, S. Farhat, A. Weckel, K. Taravati, E. G. Leitner, S. Clancy, M. Pauli, M. L. Piper, J. N. Cohen, J. F. Ashouri, M. M. Lowe, M. D. Rosenblum, T. C. Scharschmidt, Developing Human Skin Contains Lymphocytes Demonstrating a Memory Signature. Cell Reports Med. 1 (2020), doi:10.1016/j.xcrm.2020.100132.

30. M. Cichorek, M. Wachulska, A. Stasiewicz, A. Tymińska, Skin melanocytes: Biology and development. Postep. Dermatologii i Alergol. (2013), doi:10.5114/pdia.2013.33376.

31. R. L. Belote, D. Le, A. Maynard, U. E. Lang, A. Sinclair, B. K. Lohman, V. Planells-Palop, L. Baskin, A. D. Tward, S. Darmanis, R. L. Judson-Torres, Human melanocyte development and melanoma dedifferentiation at single-cell resolution. Nat. Cell Biol. (2021), doi:10.1038/s41556-021-00740-8.

32. A. Lachmann, H. Xu, J. Krishnan, S. I. Berger, A. R. Mazloom, A. Ma’ayan, ChEA: Transcription factor regulation inferred from integrating genome-wide ChIP-X experiments. Bioinformatics (2010), doi:10.1093/bioinformatics/btq466.

33. L. Domingues, I. Hurbain, F. Gilles-Marsens, J. Sirés-Campos, N. André, M. Dewulf, M. Romao, C. Viaris de Lesegno, A.-S. Macé, C. Blouin, C. Guéré, K. Vié, G. Raposo, C. Lamaze, C. Delevoye, Coupling of melanocyte signaling and mechanics by caveolae is required for human skin pigmentation. Nat. Commun. 11, 2988 (2020).

34. W. Wang, Z. Yan, J. Hu, W. J. Shen, S. Azhar, F. B. Kraemer, Scavenger receptor class B, type 1 facilitates cellular fatty acid uptake. Biochim. Biophys. Acta - Mol. Cell Biol. Lipids (2020), doi:10.1016/j.bbalip.2019.158554.

35. L. A. Kalekar, J. N. Cohen, N. Prevel, P. M. Sandoval, A. N. Mathur, J. M. Moreau, M. M. Lowe, A. Nosbaum, P. J. Wolters, A. Haemel, F. Boin, M. D. Rosenblum, Regulatory T cells in skin are uniquely poised to suppress profibrotic immune responses. Sci. Immunol. (2019), doi:10.1126/sciimmunol.aaw2910.

36. M. Ito, K. Komai, S. Mise-Omata, M. Iizuka-Koga, Y. Noguchi, T. Kondo, R. Sakai, K. Matsuo, T. Nakayama, O. Yoshie, H. Nakatsukasa, S. Chikuma, T. Shichita, A. Yoshimura, Brain regulatory T cells suppress astrogliosis and potentiate neurological recovery. Nature. 565, 246–250 (2019).

37. J. Fujisaki, J. Wu, A. L. Carlson, L. Silberstein, P. Putheti, R. Larocca, W. Gao, T. I. Saito, C. Lo Celso, H. Tsuyuzaki, T. Sato, D. Côté, M. Sykes, T. B. Strom, D. T. Scadden, C. P. Lin, In vivo imaging of T reg cells providing immune privilege to the haematopoietic stem-cell niche. Nature (2011), doi:10.1038/nature10160.

38. Y. Hirata, K. Furuhashi, H. Ishii, H. W. Li, S. Pinho, L. Ding, S. C. Robson, P. S. Frenette, J. Fujisaki, CD150 high Bone Marrow Tregs Maintain Hematopoietic Stem Cell Quiescence and Immune Privilege via Adenosine. Cell Stem Cell (2018), doi:10.1016/j.stem.2018.01.017.

39. D. Cipolletta, M. Feuerer, A. Li, N. Kamei, J. Lee, S. E. Shoelson, C. Benoist, D. Mathis, PPAR-γ is a major driver of the accumulation and phenotype of adipose tissue T reg cells. Nature (2012), doi:10.1038/nature11132.

40. A. Glasner, S. A. Rose, R. Sharma, H. Gudjonson, T. Chu, J. A. Green, S. Rampersaud, I. K. Valdez, E. S. Andretta, B. S. Dhillon, M. Schizas, S. Dikiy, A. Mendoza, W. Hu, Z.-M. Wang, O. Chaudhary, T. Xu, L. Mazutis, G. Rizzuto, A. Quintanal-Villalonga, P. Manoj, E. de Stanchina, C. M. Rudin, D. Pe’er, A. Y. Rudensky, Conserved transcriptional connectivity of regulatory T cells in the tumor microenvironment informs new combination cancer therapy strategies. Nat. Immunol. 24, 1020–1035 (2023).

41. C. Schuster, C. Vaculik, M. Prior, C. Fiala, M. Mildner, W. Eppel, G. Stingl, A. Elbe-Bürger, Phenotypic Characterization of Leukocytes in Prenatal Human Dermis. J. Invest. Dermatol. 132, 2581–2592 (2012).

42. R. A. Spritz, G. H. L. Andersen, Genetics of Vitiligo. Dermatol. Clin. (2017), doi:10.1016/j.det.2016.11.013.

43. P. Bankhead, M. B. Loughrey, J. A. Fernández, Y. Dombrowski, D. G. McArt, P. D. Dunne, S. McQuaid, R. T. Gray, L. J. Murray, H. G. Coleman, J. A. James, M. Salto-Tellez, P. W. Hamilton, QuPath: Open source software for digital pathology image analysis. Sci. Rep. 7, 16878 (2017).

