## Supplementary Figures for "Early skin seeding regulatory T cells modulate PPARγ-dependent skin pigmentation"

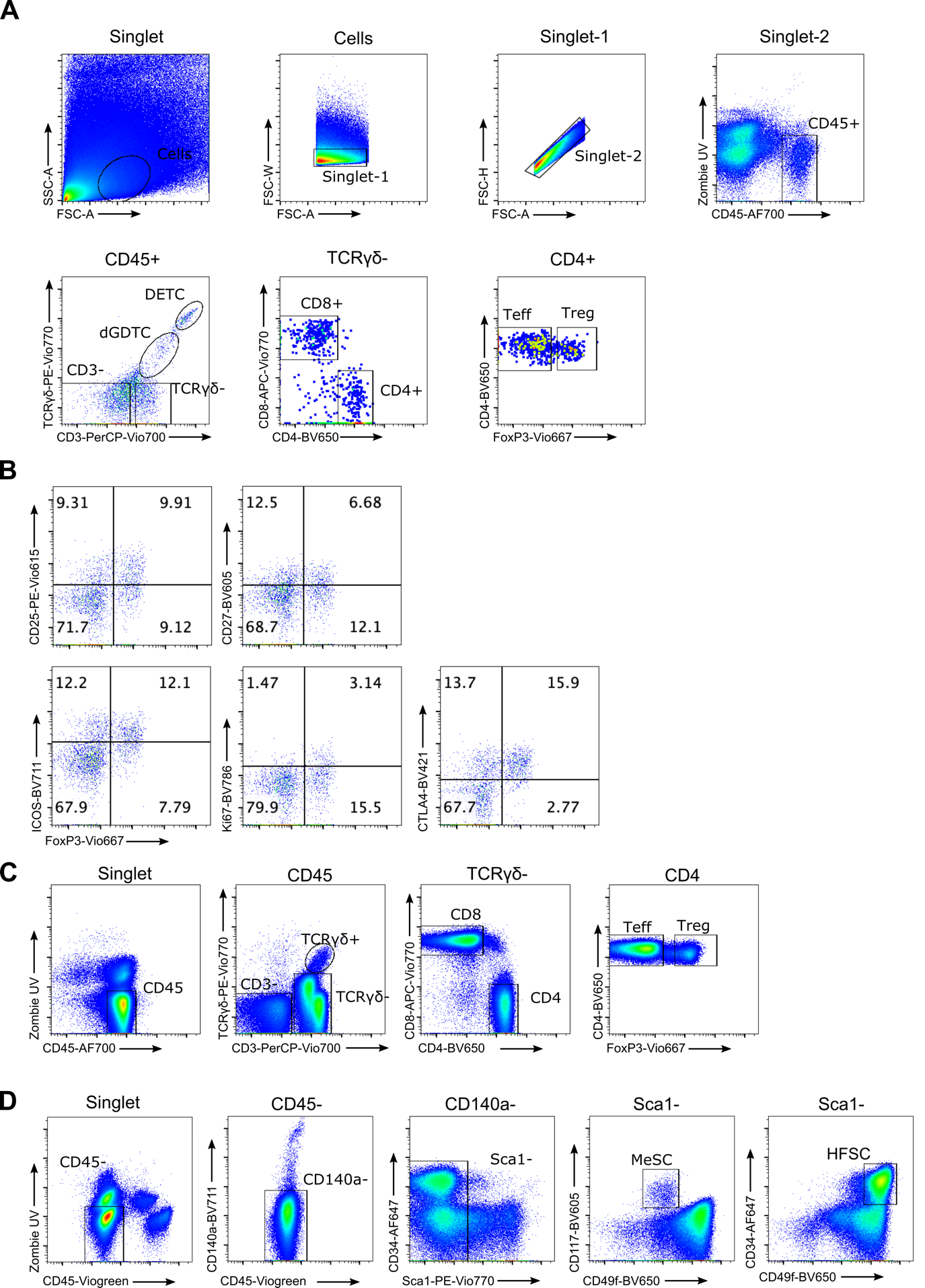

**Supplementary figure 1. Flow cytometric gating strategy for analysis**. **A)** Skin T cell gating strategy. Strict live dead and singlet inclusion criteria were applied. Immune cells were gated as CD45^+^ cells. Further subsets were gated as follows: CD3^-^ immune cells, TCRγδ^hi^ CD3^+^ DETCs (dendritic epidermal T cells), TCRγδ^int^ CD3^+^ dGDTCs (dermal gamma delta T cells), TCRγδ^-^ CD3^+^ CD8^+^ T cells, TCRγδ^-^ CD3^+^ CD4^+^ FoxP3^-^ Teffs (effector T cells), and TCRγδ^-^ CD3^+^ CD4^+^ FoxP3^+^ Tregs (regulatory T cells). **B)** Representative gating of Treg phenotypic marker expression in the skin. Cells were pre-gated as CD4^+^ T cells. **C)** Gating strategy to analyse T cells in the SDLN (skin-draining lymph node). Immune cells were gated as follows: CD3^-^ immune cells, TCRγδ^+^ cells, TCRγδ^-^ CD3^+^ CD8^+^ T cells, TCRγδ^-^ CD3^+^ CD4^+^ FoxP3^-^ Teffs (effector T cells), and TCRγδ^-^ CD3^+^ CD4^+^ FoxP3^+^ Tregs (regulatory T cells). **D)** Gating strategy to identify CD45^-^ CD140a^-^ Sca1^-^ CD117^+^ MeSCs (melanocyte stem cells) and CD45^-^ CD140a^-^ Sca1^-^ CD34^+^ CD49f^+^ hair follicle stem cells (HFSCs).

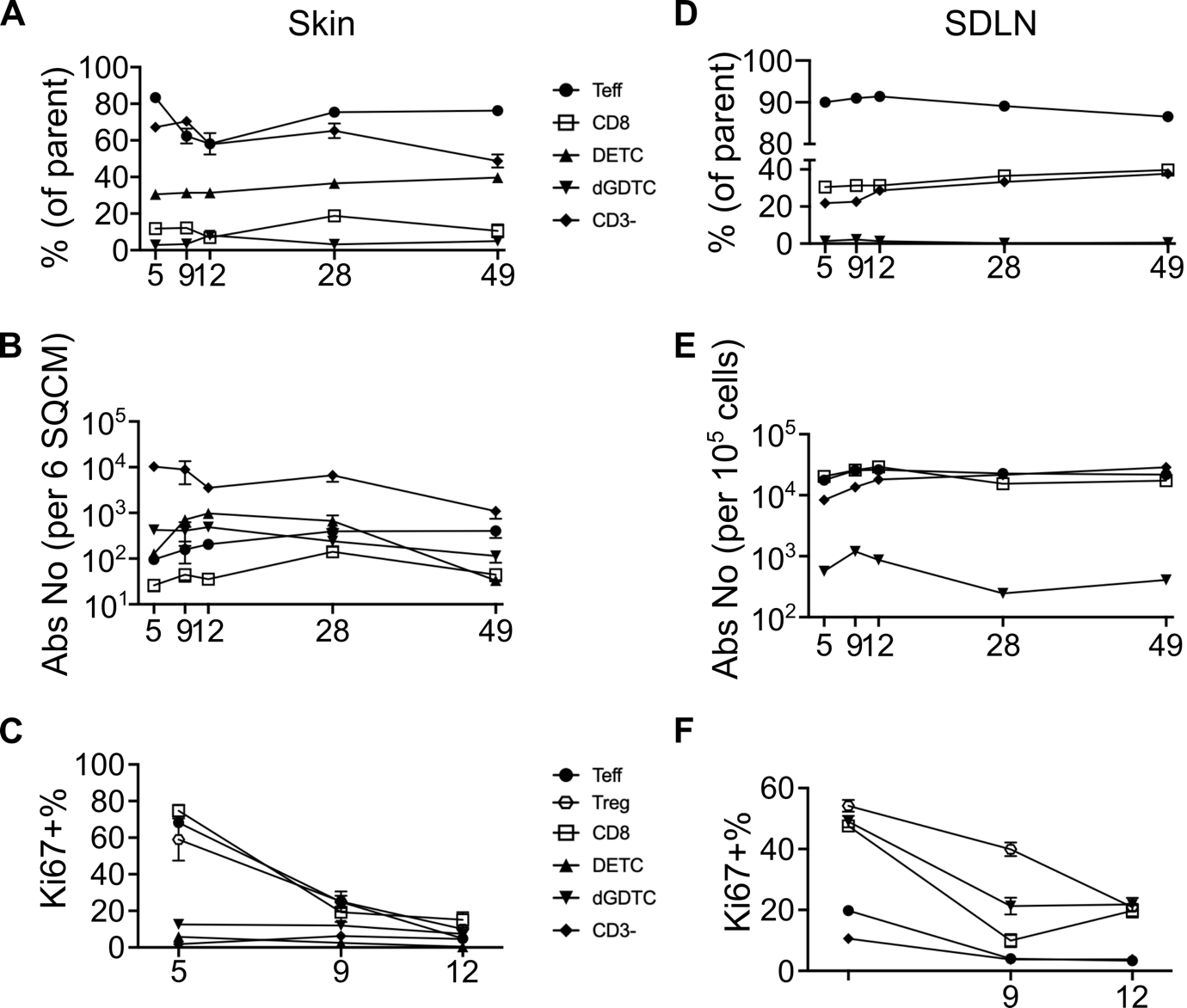

**Supplementary figure 2. Flow cytometric characterisation of T cell numbers and proliferation**. **A-C)** Skin-resident immune cells on postnatal day 5 (P5), P9, P12, P28 and P49. **A)** Percentage. **B)** Absolute number (per 6 cm^2^ of skin)**. C)** Percentage of proliferating Ki67**^+^** subsets on P5, P9 and P12. **D-F)** SDLN-resident immune cells. **D)** Percentage. **E)** Absolute number (per 10^5^ total cells). **F)** Percentage of proliferating Ki67^+^ subsets on P5, P9 and P12. Graphs show mean ± S.E.M. Data are pooled from 2 independent experiments (n=4-6 biological replicates).

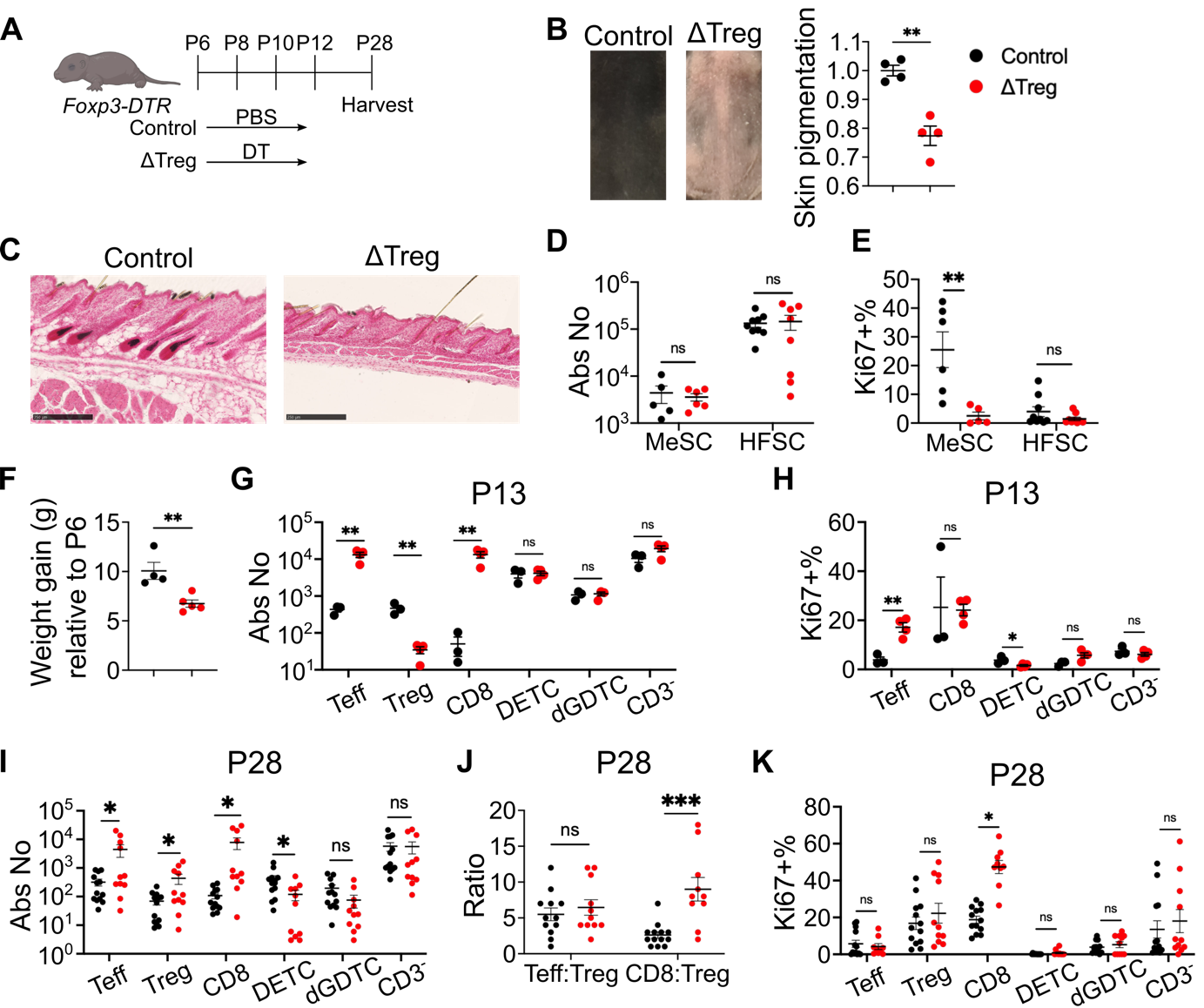
**Supplementary figure 3. Four-dose DT regimen causes weight loss, inflammation, and pigment defect. A)** Schematic outline of experimental timeline. Tregs were depleted by intraperitoneal administration of diphtheria toxin (DT) on postnatal day 6 (P6), P8, P10 and P12 (30 ng/g per injection). Skin tissues were harvested on P28. **B)** Representative images of dorsal skin on P28, and associated quantification of skin pigmentation. **C)** Fontana & Masson (F&M) staining of P28 dorsal skin. Scale bar represents 250 µm. **D-E**) Flow cytometric quantification of melanocyte stem cells (MeSCs) and hair follicle stem cells (HFSCs). **D)** Absolute number. **E)** Percentage of proliferating Ki67^+^ cells. **F**) Weight gain from P28 to P6. **G-H)** Flow cytometric quantification of skin T cells on P13. **G)** Absolute number. **H)** Percentage of proliferating Ki67^+^ cells. **I-K)** Flow cytometric quantification of skin T cells on P28. **I)** Absolute number. **J)** Ratio of Teff:Treg and CD8:Treg. **K)** Percentage of proliferating Ki67^+^ subsets. Data are pooled from 4 independent experiments (n=2-4 biological replicates). Graphs show mean ± S.E.M. Unpaired t-test. ***p<0.001, **p<0.01, *p<0.05, ns p>0.05.

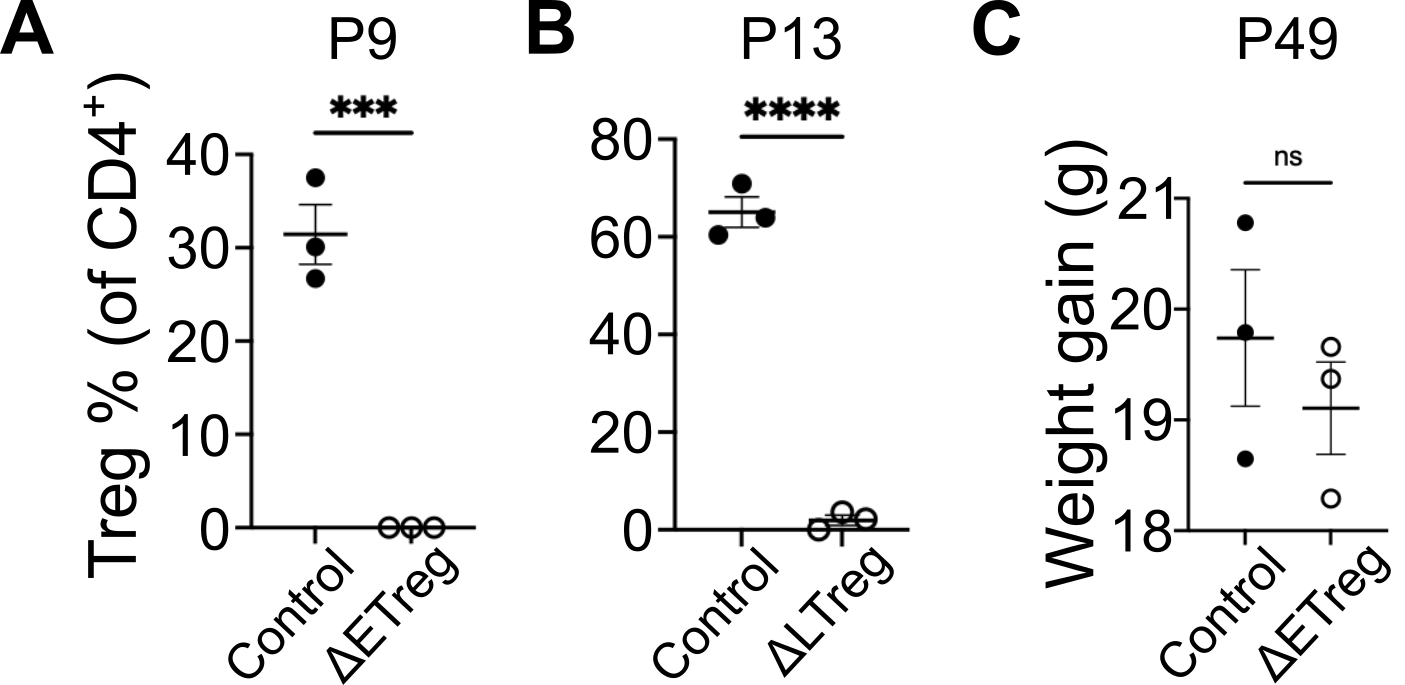
**Supplementary figure 4. Transient DT injection is sufficient for skin Treg depletion.** Flow cytometric profiling of Treg percentage on **A)** P9 or **B)** P13 following Treg depletion in *Foxp3-DTR* mice. **C)** Weight gain from P6 to P49 in grammes. Graphs show mean ± S.E.M. Unpaired t-test. ****p<0.0001, ***p<0.001, ns p>0.05.

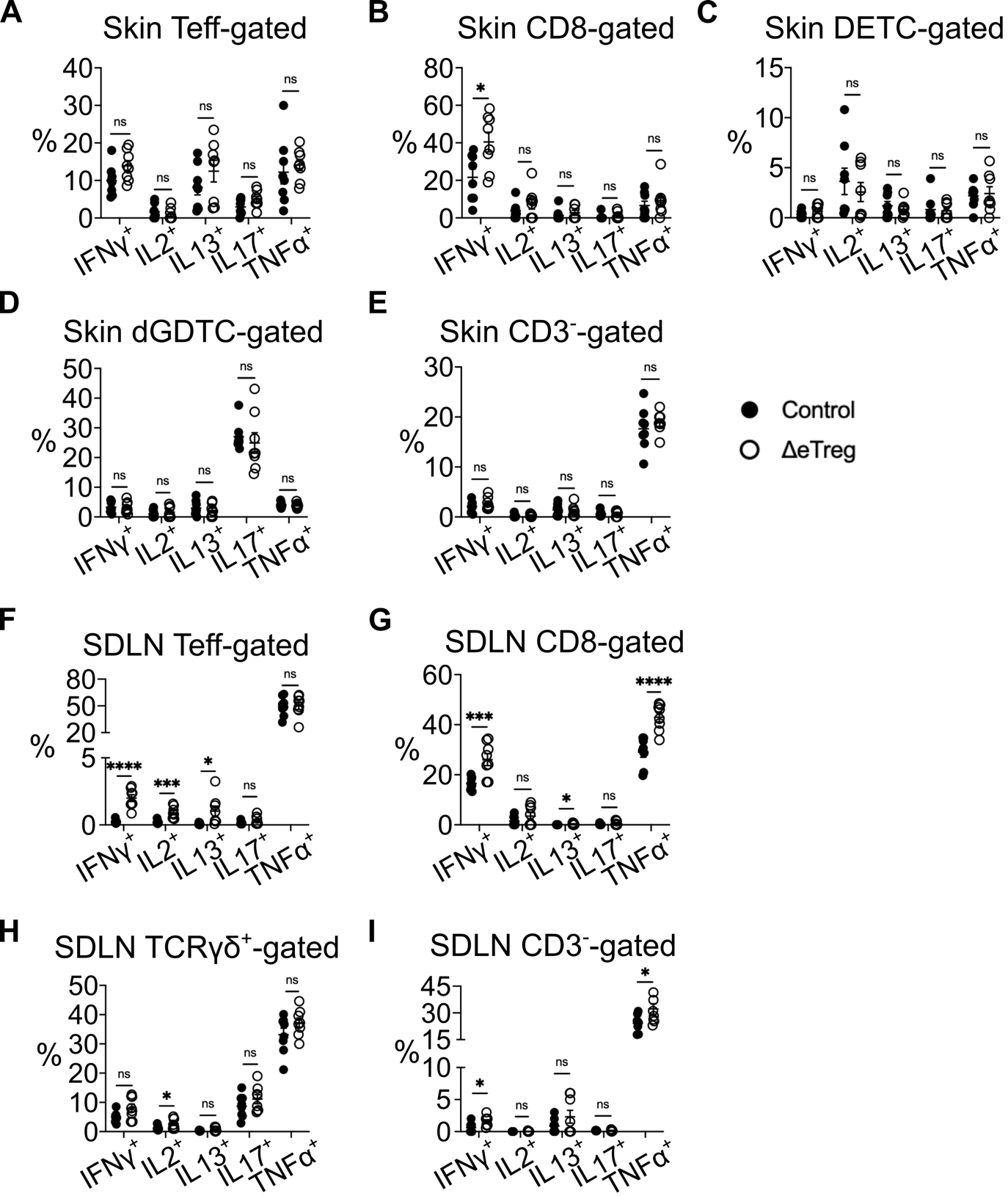
**Supplementary figure 5. Early Treg depletion causes inflammatory response in the skin and SDLN.** Percentage production of cytokines interferon-γ (IFNγ), interleukin 2 (IL2), IL13, IL17 and tumour necrosis factor α (TNFα) on P9. **A-I)** Percentage expression of cytokines following stimulation with PMA/Ionomycin. **A-E)** Skin. **A)** Teffs. **B)** CD8^+^ T cells. **C)** DETCs. **D)** dGDTCs. **E)** CD3^-^ immune cells. **F-I)** SDLN. **F)** Teffs. **G)** CD8^+^ T cells. **H)** TCRγδ^+^ T cells. **I)** CD3^-^ immune cells. Data are pooled from two independent experiments. Graphs show mean ± S.E.M (n=4 biological replicates). Unpaired t-test. ****p<0.0001, ***p<0.001, *p<0.05, ns p>0.05.

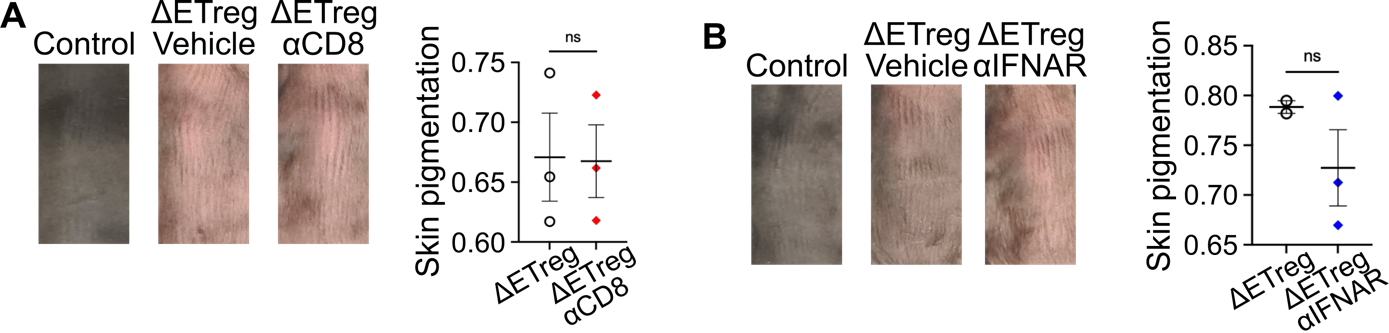
**Supplementary figure 6. Modulation of skin pigmentation. A-D)** Mice were intraperitoneally injected on P6 and P8. Dorsal skin tissues were harvested and photographed on P28. **A)** *Foxp3-DTR* mice received PBS (control) or DT (ΔEtreg) in addition to PBS (vehicle) or 100 µg of α-CD8 depleting antibody (αCD8) per injection. αCD8 groups received further injections on P15 and P22. **B)** *Foxp3-DTR* mice received PBS (control), or DT (ΔEtreg) in addition to PBS (vehicle) or 100 µg of α-IFNAR blocking antibody (αIFNAR) per injection. Graphs show mean ± S.E.M. Data shows biological replicates (n=2-3). Unpaired t-test. ns p>0.05.

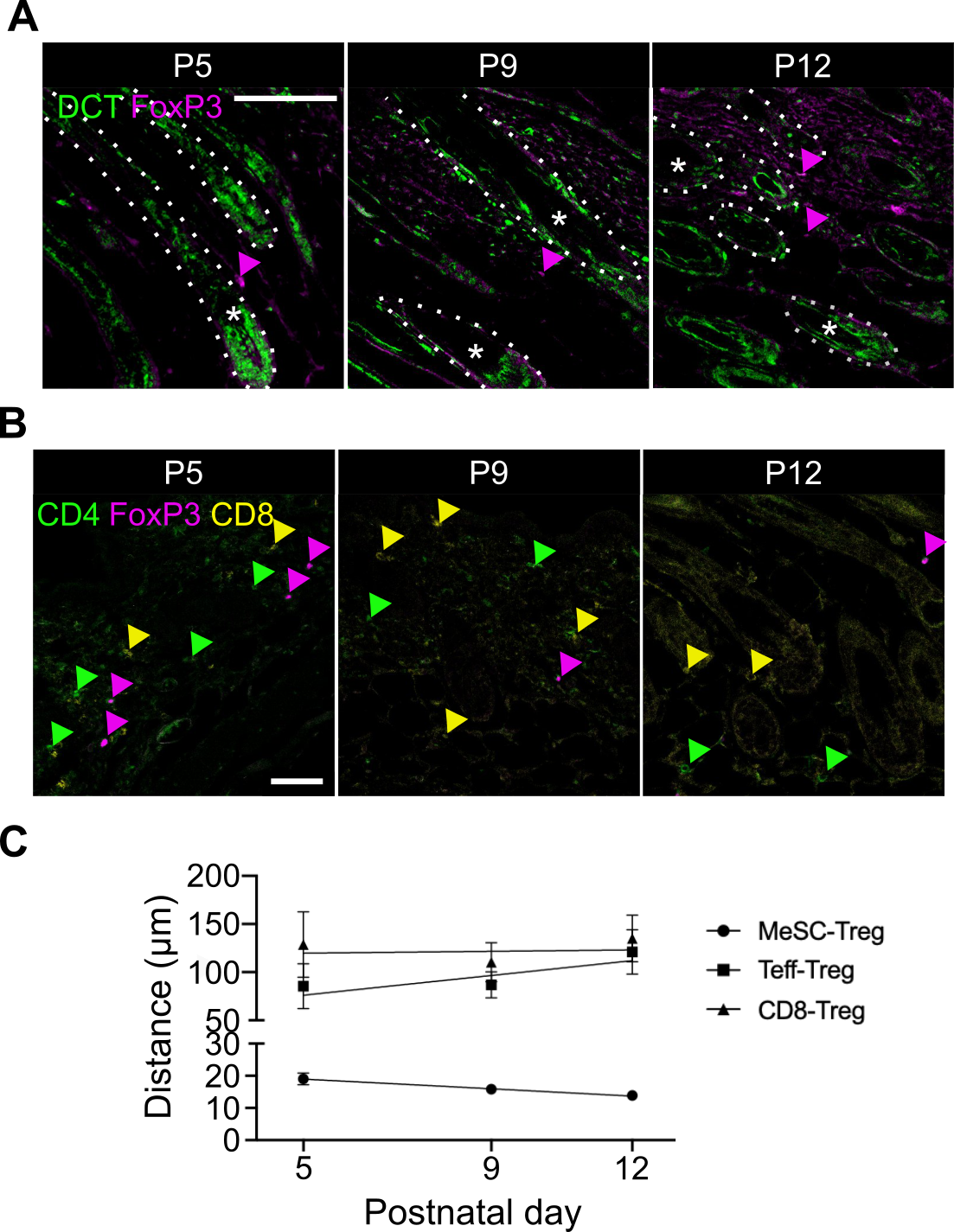
**Supplementary figure 7. Skin Tregs reside proximally to melanocyte stem cells, and not T cells. A-B)** Histological quantification of Treg distance to melanocyte stem cells (MeSCs) on postnatal day 5 (P5), P9 and P12. **A)** Immunofluorescence staining of DCT^+^ MeSCs and Foxp3^+^ Tregs. Dotted lines demarcate hair follicles. **B)** Immunofluorescence staining of CD4^+^ FoxP3^-^ effector T cells (Teff, green arrowhead), CD4^+^ FoxP3^+^ Tregs (Magenta arrowhead), and CD8^+^ cytotoxic T cells (CD8, yellow arrowhead). **C)** Quantification of MeSC-Treg, Teff-Treg and CD8-Treg distances on P5, P9 and P12. Graph shows mean ± S.E.M. Data are pooled from 3 independent experiments (n=3-6 biological replicates).

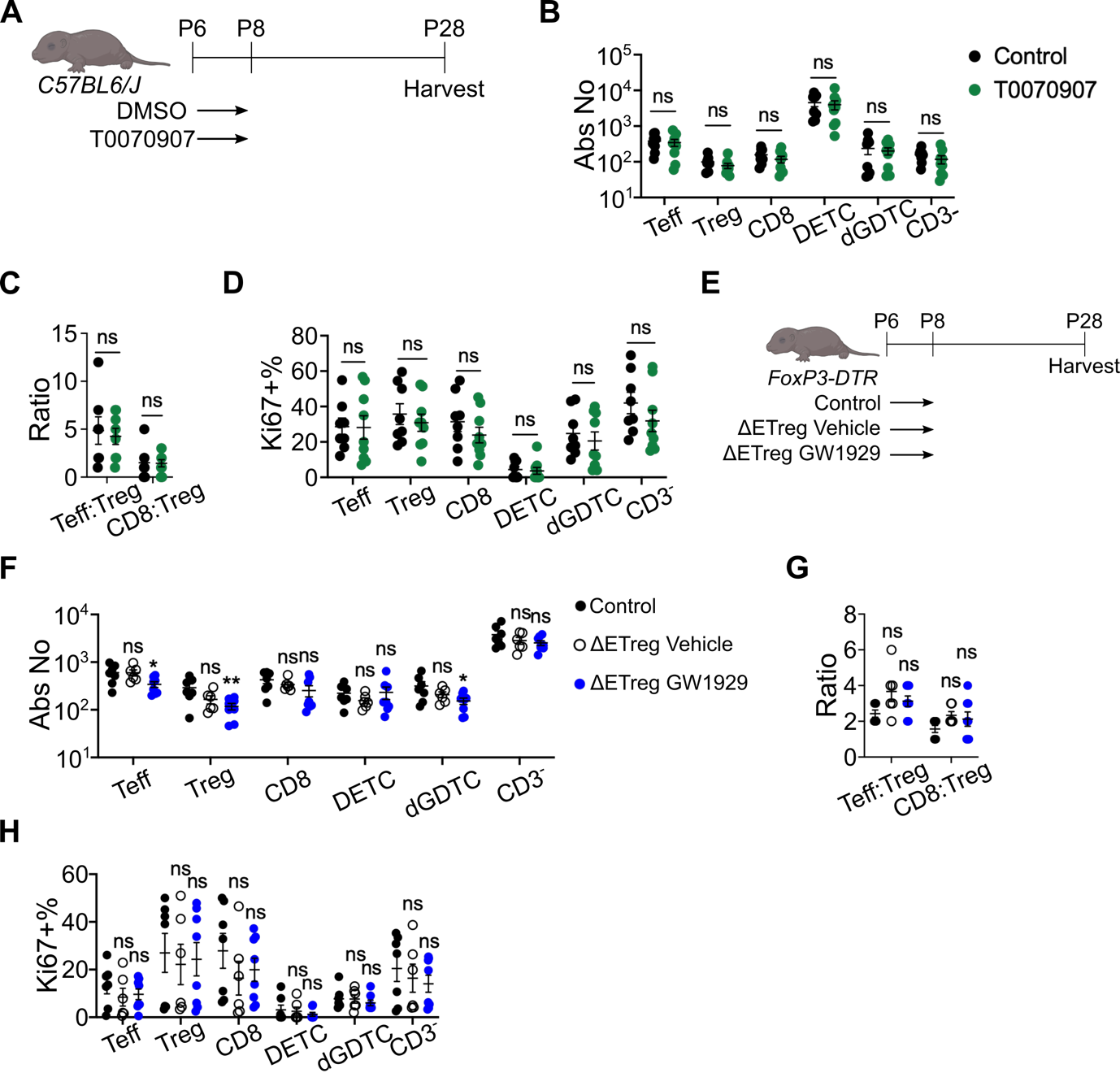
**Supplementary figure 8. Modulation of neonatal PPARγ signalling does not affect skin-resident immune cell numbers or proliferation. A)** Schematic of experimental timeline. C57BL6/J mice received 2.5 µg/g of T0070907 or DMSO on P6 and P8. Skin tissues were harvested on P28. **B-E)** Flow cytometric quantification of skin-resident T cells and Tregs. **B)** Absolute number. **C)** Teff:Treg and CD8:Treg ratio. **D)** Percentage of proliferating Ki67^+^ cells. **E)** Schematic outline of rescue experiment using PPARγ agonist. Control group received PBS and DMSO on P6 and P8. Τreg-depleted ΔEtreg groups received DMSO (vehicle) or GW1929 in addition to DT on P6 and P8. Skin tissues were harvested on P28. **F)** Absolute number. **G)** Teff:Treg and CD8:Treg ratio. **H)** Percentage of proliferating Ki67^+^ cells. Data are pooled from two independent experiments. Graphs show mean ± S.E.M. (n=4-5 biological replicates). Unpaired t-test. ns p>0.05.
